# MATES: A Deep Learning-Based Model for Locus-specific Quantification of Transposable Elements in Single Cell

**DOI:** 10.1101/2024.01.09.574909

**Authors:** Ruohan Wang, Yumin Zheng, Zijian Zhang, Kailu Song, Erxi Wu, Xiaopeng Zhu, Tao P. Wu, Jun Ding

## Abstract

Transposable elements (TEs) are crucial for genetic diversity and gene regulation. Current single-cell quantification methods often align multi-mapping reads to either ‘best-mapped’ or ‘random-mapped’ locations and categorize them at subfamily levels, overlooking the biological necessity for accurate, locus-specific TE quantification. Moreover, these existing methods are primarily designed for and focused on transcriptomics data, which restricts their adaptability to single-cell data of other modalities. To address these challenges, here we introduce MATES, a deep-learning approach that accurately allocates multi-mapping reads to specific loci of TEs, utilizing context from adjacent read alignments flanking the TE locus. When applied to diverse single-cell omics datasets, MATES shows improved performance over existing methods, enhancing the accuracy of TE quantification and aiding in the identification of marker TEs for identified cell populations. This development facilitates the exploration of single-cell heterogeneity and gene regulation through the lens of TEs, offering an effective transposon quantification tool for the single-cell genomics community.

## Introduction

Transposable elements (TEs), also known as transposons or jumping genes, form a significant part of mammalian genomes and play crucial roles in gene regulation, genome evolution, and cell-to-cell heterogeneity [1–3]. Although some TEs are still active and jump in our genome, the majority have accumulated mutations and degenerations that render them incapable of active transposing. Consequently, many TEs are retained in the genome and serve as regulatory elements. These non-coding functions include the regulation of gene expression and the formation of long non-coding RNAs (lncRNAs), which are involved in critical regulatory networks that influence gene expression and cellular function [1, 4]. Despite these important roles, our understanding of locus-specific TEs at the single-cell level has been limited due to challenges in quantifying multi-mapping sequencing reads, a result of their repetitive sequences and high copy numbers [5].

Studying TEs in single-cell genomics, in contrast to bulk sequencing, is important for understanding their dynamic regulation and contribution to cellular heterogeneity. This approach reveals the complex expression patterns of TEs and their significant impact on the transcriptional landscape [6, 7]. The variability in TE activity across individual cells contributes to the complexity of gene regulation and cellular dynamics, crucial in both normal development and disease states like cancer [8, 9].

While there is an abundance of methods for TE quantification at the bulk level, such as Squire [10], TEtranscripts [11], and Telescope [12], emerging single-cell methods like scTE [2] and SoloTE [13] remain limited. SoloTE addresses the challenge of repetitive sequence assignment by allocating multi-mapping reads to a singular locus based on the best read alignment score. In contrast, scTE employs a similar strategy but rectifies multi-mapping errors using the count of unique-mapping reads. However, these methods often rely heavily on alignment algorithms to address multi-mapping reads, neglecting the genomic context flanking the TEs. Furthermore, current methods fail to provide an accurate locus-specific TE quantification. scTE measures TE expression by allocating reads to corresponding TE subfamilies. SoloTE did present locus-level TE expression, but it only maps the reads to the “best” location; thus, it is very limited in handling multi-mapping reads that prevail in TE regions [14, 15]. These approaches overlook or circumvent the challenging multi-mapping read assignment rooted in the repetitive characteristics of TEs. This oversight can result in an underestimation of the complexity and uncertainty associated with assigning multi-mapping reads in TE quantification. The frequent occurrence of multi-mapping reads across TE regions highlights the existing gaps in single-cell TE quantification techniques, emphasizing the need for precise read assignment to specific genomic loci. An accurate quantification of locus-specific TE expression is paramount to understanding the mechanisms of TEs’ crosstalk with regional genome regulations [6, 16].

Recent advancements in single-cell sequencing technologies have broadened the scope to include a range of modalities [3, 17, 18], extending beyond traditional transcriptomics. These developments allow for the analysis of different cellular components, such as the epigenome, in addition to the transcriptome, within individual cells. Furthering this progression, single-cell multi-omics methods, like 10x Genomics’ Multiome, can even simultaneously profile both the transcriptome and epigenome in the same cells. This dual-modality profiling is highlighted in comprehensive reviews of the technological landscape and applications in single-cell multi-omics, demonstrating a significant impact on molecular cell biology and the potential to decipher complex biological processes at the single-cell level [19–22]. Despite these advancements, current TE quantification methods, primarily designed for single-cell transcriptomics, face limitations when dealing with data from other modalities, such as single-cell assays for transposase-accessible chromatin sequencing (scATAC-seq). These methods also lack comprehensive solutions for the combined quantification and analysis of TEs in multi-omics datasets. Consequently, there is a pressing need for methodologies capable of accurately aligning multi-mapping TE reads and performing locus-level quantification across various modalities, underscoring the ongoing challenges and opportunities in the field of single-cell genomics and multi-omics research, particularly through the evolving lens of TEs. These advancements must also extend to non-mammalian species, where TE dynamics play a crucial role in understanding broader biological phenomena [23, 24].

To address these challenges and fill the gap, we introduce MATES (Multi-mapping Alignment for TE loci quantification in Single-cell), a deep neural network-based method tailored for accurate locus-specific TE quantification in single-cell sequencing data across modalities. MATES harnesses the distribution of uniquely mapped reads occurrence flanking TE loci and assigns multi-mapping TE reads for locus-specific TE quantification. By leveraging the power of a deep neural network, MATES captures complex relationships between the distribution of unique-mapping reads around TE loci and the probability of multi-mapping reads being assigned to those loci. This approach allows MATES to handle multi-mapping read assignments probabilistically, based on the local context surrounding the TE loci.

In our systematic evaluation of MATES using various single-cell datasets across different sequencing platforms, modalities and species, we demonstrated that MATES consistently provides more accurate TE quantification compared to existing methods. In addition to higher precision, MATES offers locus-specific TE quantification and can be generalized across different sequencing platforms and data modalities, enabling a more comprehensive understanding of how TEs contribute to cellular dynamics and gene regulation. We also validated the method’s predictions using nanopore and PacBio long-read sequencing, as well as simulations. By comparing MATES’s single-cell TE quantification with the ground truth from simulations or the proxy ground truth from long-read sequencing, we demonstrated MATES’s accuracy and its advantage over current methods. Our results indicate that MATES is effective for exploring the roles of TEs in single-cell biology and provides a practical solution for TE quantification in various experimental contexts. MATES addresses the existing gap in single-cell TE quantification techniques, providing a robust and adaptable method that improves our ability to understand the contributions of TEs to cellular heterogeneity and genome regulation. As the field of single-cell sequencing continues to grow, the potential for in-depth TE quantification and analysis expands, offering new opportunities to gain insights into the molecular mechanisms underlying various biological processes [9].

## Results

### Methods overview

MATES is a specialized tool designed for locus-level quantification of TEs in single-cell datasets of different modalities. The method involves several key steps. First, raw reads are mapped to the reference genome, identifying reads that map uniquely to a TE locus (unique reads) and reads that map to multiple TE loci (multi-mapping reads) (Fig.1 a). Next, we compute the coverage vector for each TE locus, representing the distribution of unique reads surrounding the locus (context). Each TE region (locus) is then subdivided into smaller bins of length *W* (e.g., 10 base pairs). Each of these bins is classified into Unique-dominant (U) or Multi-dominant regions (M) based on the percentage of unique and multi-mapping reads within the bin (Fig.1 b). Please refer to the Methods section for details on selecting these hyper-parameters. Third, we employed an AutoEncoder (AE) model to learn the latent embeddings (*V_u_*) that represent the high-dimensional unique reads coverage vector of a TE locus, which indicates the mapping context flanking the specific TE locus. The one-hot encoded TE family information (*T_i_*) was also taken as input for the model. Fourth, the learned latent embedding (*V_u_*) and the TE family embedding (*T_i_*) are used to predict the multi-mapping ratio (*α*) for the specific TE locus via a multiple-layer perceptron regressor. The grand used loss to learn the model is composed of two components (*L*_1_ and *L*_2_). The former represents the reconstruction loss of the AutoEncoder, while the latter reflects the continuity of the actual reads coverage across neighboring small bins on the TE. Essentially, the final read coverage on the multi-mapping dominant (M) bins should be close to its neighboring unique dominant (U) bins because of their genomics proximity. Finally, once we train the model that predicts the multi-mapping ratio for each TE locus, we can leverage it to count the total number of reads that fall into the specific TE loci, presenting probabilistic quantification of TEs at the locus level (Fig.1 c). By combining TE quantification with conventional gene quantification (e.g., gene expression or gene accessibility) from the single-cell data, which we refer to as “Gene+TE expression” in the sections below, we can then more accurately cluster the cells and identify comprehensive biomarkers (genes and TEs) to characterize the obtained cell clusters (cell sub-populations). Equipped with advanced features, MATES efficiently handles various single-cell data modalities. Its application offers insights into the roles of TEs in diverse datasets, clustering cells, and identifying potential biomarker TEs (Fig.1 d). Beyond its analytical capabilities, MATES also presents locus-specific TE visualization and interpretation. The tool facilitates the generation of comprehensive bigwig files and Interactive Genomic Viewer (IGV) plots, enabling researchers to visually explore and interpret read assignments for TE loci across the genome (Fig.1 e). This capability unlocks the investigation of potential interactions between TEs and the genes situated near TE loci, significantly enhancing our understanding of TE dynamics and their impact on gene regulation and cellular functions. Please note that, except where specifically mentioned, the term ‘TE’ used throughout this manuscript represents repetitive elements identified from RepeatMasker. This allows us to provide a comprehensive overview of genomic repeats in our study. When discussing the ‘stricter’ definition of TEs, we have specifically mentioned which TE subfamilies are included.

**Fig. 1.**
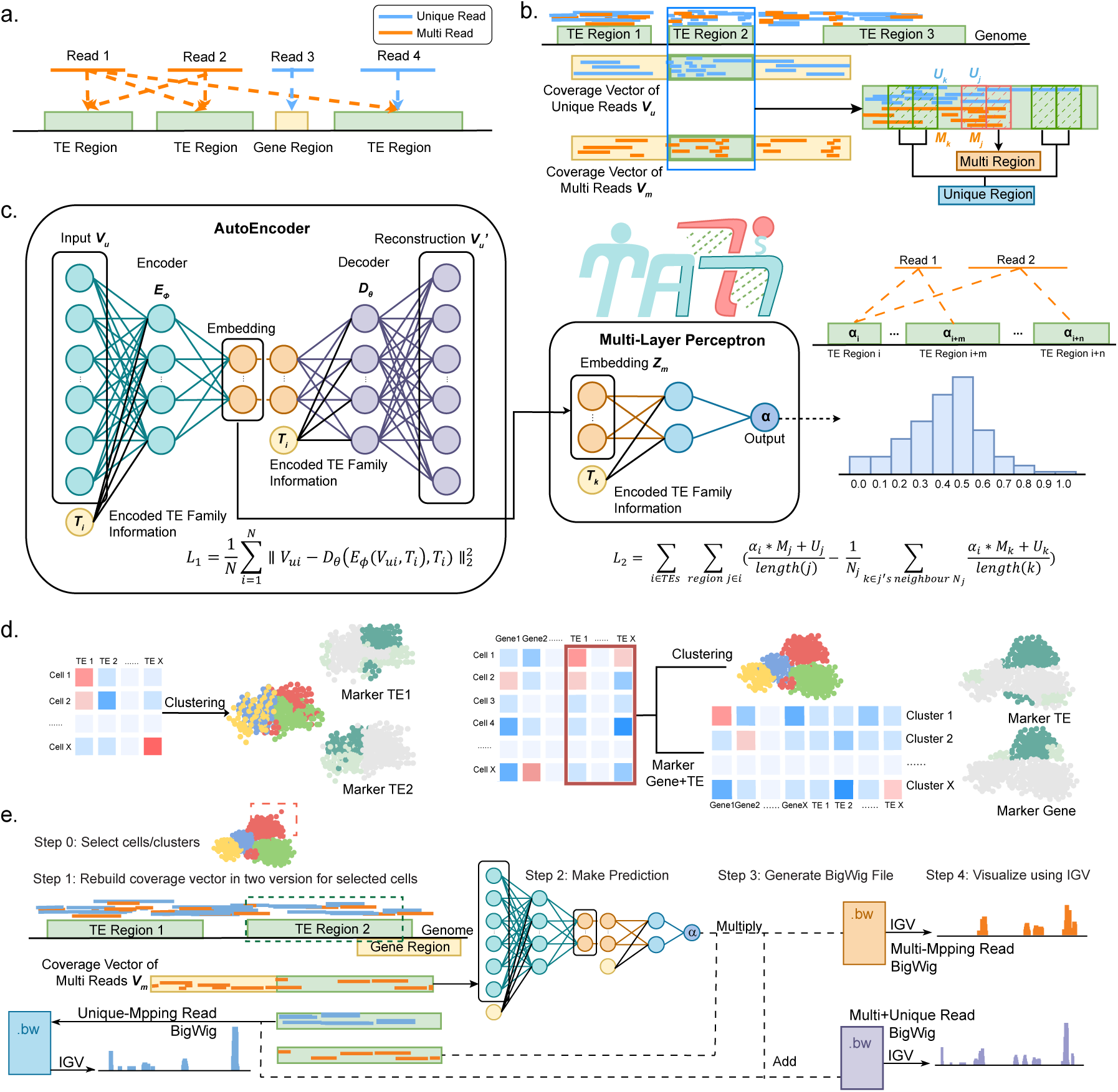
MATES methodology for TE quantification and analysis. (a) Raw reads are aligned to the reference genome, accounting for multi-mapping reads at TEs’ loci. (b) TE coverage vectors, including unique reads coverage vector *V_u_* and multi reads coverage vector *V_m_*, are constructed, capturing reads’ distribution information. (c) An AutoEncoder model extracts latent embeddings from unique reads coverage vectors. These embeddings, combined with TE family data *T_i_*, predict the likelihood, *α*, of multi-mapping reads aligning to each TE locus. (d) The multi-mapping probability *α*, computed by MATES, is critical in creating the TE count matrix. This matrix is pivotal for cell analyses and can be utilized either independently or in conjunction with a conventional gene count matrix. Such combined use enhances cell clustering and biomarker (gene and TE) discovery, providing a more comprehensive understanding of cellular characteristics. (e) Genome-wide reads coverage visualization by MATES in the Genome Browser. This method quantifies TEs at specific loci in individual cells, producing bigwig files with coverage from probabilistically assigned multi-mapping reads. These files, containing both unique and multi-mapping reads, are merged to generate comprehensive bigwig files for genome-wide TE read visualization using tools such as the Interactive Genomic Viewer (IGV).

### MATES identifies signature TEs and their specific loci in 2C-like cells (2CLCs) within 10x single-cell RNA-seq data of chemical reprogramming

To demonstrate the precision of MATES in TE quantification from single-cell RNA-seq data, we applied it to a 10x single-cell chemical reprogramming dataset of mice. This analysis identified signature TEs of 2-cell-like cells (2CLCs) [25]. By employing MATES to quantify TE expressions, we integrated the quantified TE count matrix with gene expression profiles, facilitating comprehensive clustering and visualization analyses, as shown in Fig.2 a,b and Supplementary Fig.S1 a. Our study revealed a distinct subpopulation of 2CLCs (cluster 17), positioned between stage II and stage III of reprogramming. Notably, MATES detected the 2CLCs population and distinguished their signature gene markers, especially *Zscan*4*d* and *Zscan*4*c* [26], within the transition-stage clusters. Moreover, MATES identified specific TE markers, MERVL-int and MT2_Mm, that are enriched within the 2CLC cluster, corroborating previous studies recognizing these TEs as defining markers for 2CLCs [27–30].

**Fig. 2.**
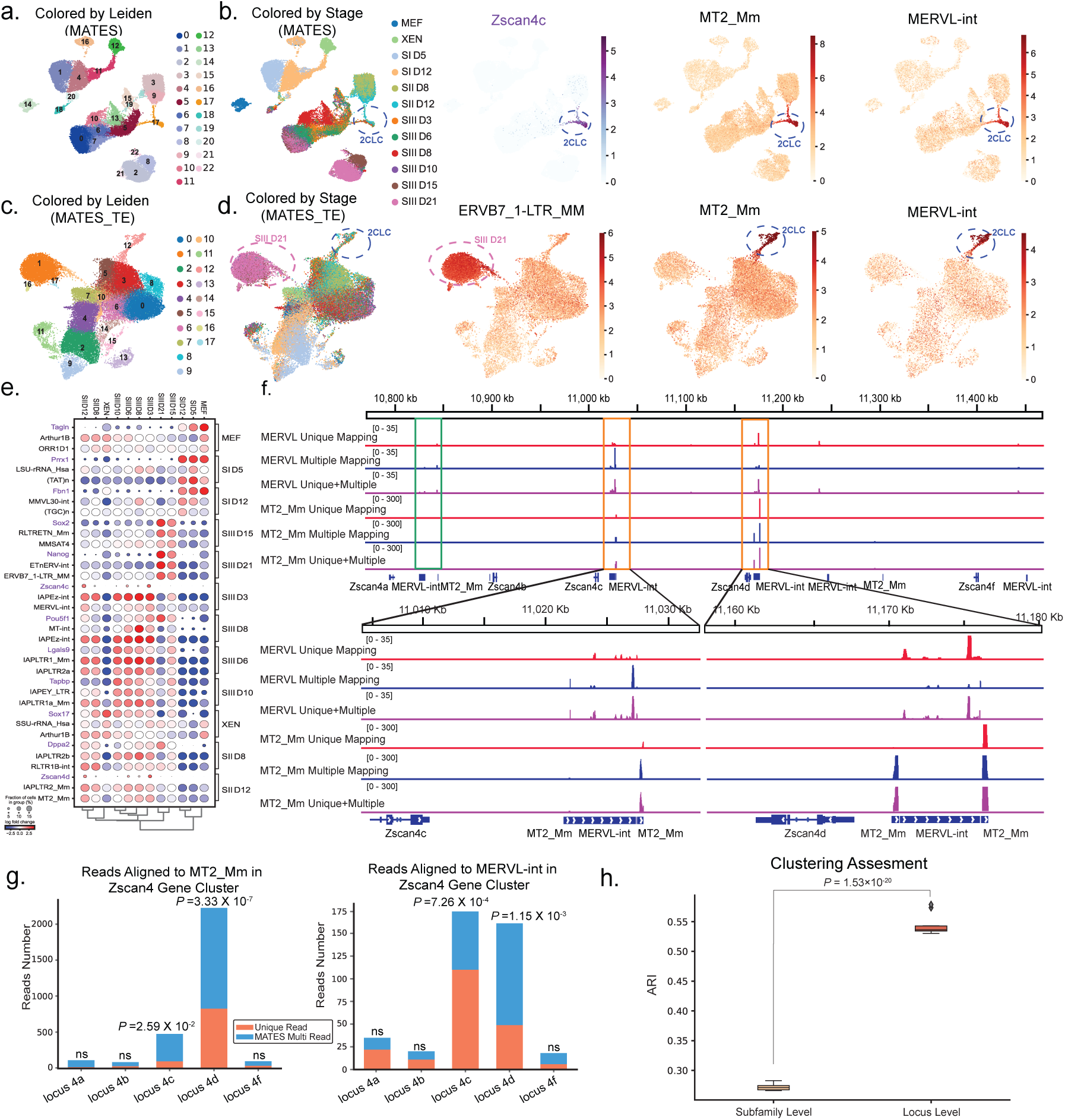
MATES enhances cell clustering and biomarker discovery in mouse chemical reprogramming. (a-b) UMAP plots illustrating MATES’s efficacy in cell clustering by integrating TEs and genes. (a) is colored by Leiden clustering results, while (b) is colored according to reprogramming stages, highlighting identified gene (purple) and TE (red) biomarkers. (c-d) Additional UMAP plots emphasizing MATES’s capability for clustering using only TEs, with (c) colored by Leiden clusters and (d) by reprogramming stages. Notably, MT2_Mm and MERVL-int TEs are prominent biomarkers in *Zscan*4*c*/*Zscan*4*d*-positive cells, consistent with known 2CLCs markers. (e) Dot plot identifying stage-specific marker genes (purple) and TEs (black) as detected by MATES. (f) Illustration of MATES’s probabilistic approach to allocating multi-mapping reads to specific TE loci, predominantly involving MT2_Mm and MERVL-int at the *Zscan*4*c*/*Zscan*4*d* loci in 2CLCs. (g) Bar plots displaying the reads enrichment for MT2_Mm and MERVL-int at the *Zscan*4*c*/*Zscan*4*d* loci. The enrichment p-value is calculated with a binomial test. (h) Box plot comparison of cell clustering efficacy by Adjusted Rand Index (ARI) between MATES’s locus-level and subfamily-level TE quantification. The experiments run with N=10 different seeds and the p-value was calculated using a one-sided Student’s t-test.

These findings highlight MATES’s ability to capture cell populations and their significant biological markers (genes and TEs), providing insights into the cellular dynamics of reprogramming.

We next conducted a TE-centric analysis to further validate the distinct role of MATES’s TE expression quantification in cell clustering and biomarker discovery (Fig.2 c,d, Supplementary Fig.S1 d). When quantifying TE expression, we took care to exclude overlapping regions between TEs and their adjacent genes to prevent potential information leakage from gene expression data. This TE-centric analysis specifically focused on TE expressions and it also identified the 2CLC cell population. Furthermore, this analysis not only confirmed the previous findings related to the 2CLCs population but also reaffirmed the relevance of its associated TE biomarkers, namely MERVL-int and MT2_Mm, as illustrated in Fig.2 c,d. This indicates that our cell clustering and biomarker discovery are not solely dependent on traditional gene expression analysis. Instead, TE quantification independently conducted by MATES provides consistent cell clustering results and accurately identifies signature TEs for the identified cell populations. To provide a clearer, quantitative view of the clustering accuracy based solely on TEs, we included confusion matrices and calculated Adjusted Rand Index (ARI) and Normalized Mutual Information (NMI) scores to highlight the similarity between TE-based and conventional gene-based analysis results. The clustering results based on TE expression alone were compared to those based on gene expression. Major clusters, such as cluster 1 and cluster 12, representing SIII D12 and 2CLCs respectively, were effectively captured by TE-only clusters (Supplementary Fig.S2 a). These TE clusters corresponded well with gene expression clusters, with high ARI (median 0.397, *P <* 1 × 10*^−^*^6^) and NMI (median 0.496, *P <* 1 × 10*^−^*^6^) scores indicating a strong alignment (Supplementary Fig.S2 b). The confusion matrix in Supplementary Fig.S2 c shows that TE cluster 1 is predominantly identified by gene clusters 1 and 9, while TE cluster 12 is mainly identified by gene cluster 18, highlighting a notable correspondence between TE and gene expression clusters. Moreover, by focusing on clustering driven by TE expression, quantified exclusively from multi-mapping reads, MATES demonstrates its ability to manage these challenging reads and identify biomarkers precisely aligned with their specific developmental stages (Supplementary Fig.S1 g).

Not only does MATES identify the signature genes and TE markers for 2CLCs and cell populations in different reprogramming stages (Fig.2 e, Supplementary Fig.S1 h), but it also shows effectiveness in aligning the multi-mapping reads to specific loci, a challenge that has stymied current methods. For instance, scTE is limited to assigning multi-mapping reads to the metagene (TEs of the same subfamily), lacking a distinct assignment to specific genomic loci. While SoloTE quantifies reads uniquely mapped to TEs at the locus level, it retains only the best alignment for multi-mapping reads and then quantifies them at the subfamily level. In contrast, by leveraging the learned multi-mapping rate for each TE locus (*α*), MATES probabilistically assigns multi-mapping reads to TE genomic loci across the genome. Through this strategy, we can quantify TE expression at the locus level with accuracy, evident when assessing the multi-mapping reads for 2CLC cells (Fig.2 f,g). Multi-mapping reads linked to MT2_Mm and MERVL-int were observed to align closely with genes *Zscan*4*c* and *Zscan*4*d*, and the total reads linked to MT2_Mm and MERVL-int that aligned closely with *Zscan*4*c* and *Zscan*4*d* loci were significantly higher than other control loci (Fig.2 g, Supplementary Fig.S1 i). This alignment is consistent with findings by Zhu et al. [31], where the activation of *Zscan*4*c* was correlated with the activation of the endogenous retrovirus MT2/MERVL. Please note that the unique reads-based locus quantification, highlighted in orange in panel g, represents the SoloTE strategy. This strategy processes unique reads at the locus level and multi-mapping reads at the subfamily level. Thus, at the locus level, only unique reads were leveraged by SoloTE, which may result in missing reads mapped to critical loci such as 4c and 4d, indicating its potential limitations. Additionally, locus-specific TE quantification improves clustering accuracy compared to the subfamily level TE quantification typically used in existing methods, clearly demonstrated in Fig.2 h. This emphasizes the substantial benefits of precise, locus-level TE quantification. For additional results demonstrating the effectiveness of MATES with this 10x single-cell RNA-seq data, please see Supplementary Fig.S1.

### MATES quantifies disease-related TE expression in full-length Smart-Seq2 single-cell RNA-seq data of Human Glioblastoma

To demonstrate the cross-platform applicability of MATES, we tested and applied the tool to another single-cell RNA-seq dataset from the Smart-Seq2 full-length sequencing platform [32], focusing on a human glioblastoma dataset [33]. The combined use of MATES’s TE expression quantification and conventional gene expression analysis allowed us to pinpoint distinct cell populations within the glioblastoma microenvironment, as shown in the UMAP plots (Fig.3 a,b). We observed certain TEs with expression patterns linked to crucial glioma gene markers like *EGFR* [33, 34] and TE markers including HUERS-P1-int [35], HERVK-int [36], along with immune cell gene markers such as *CD*74 [37, 38] and TE marker LTR2B [39, 40] (Fig.3 b). These correlations suggest that TEs might be associated with processes related to tumor heterogeneity and the immune response in glioblastoma. Further research is necessary to explore any causal relationships and the underlying mechanisms. Combining TE-based cell typing with gene expression data revealed a detailed interplay between genes and TEs. This integration showcased how TE-based clustering could complement gene expression analysis, thus enhancing the resolution of cellular heterogeneity studies.

**Fig. 3.**
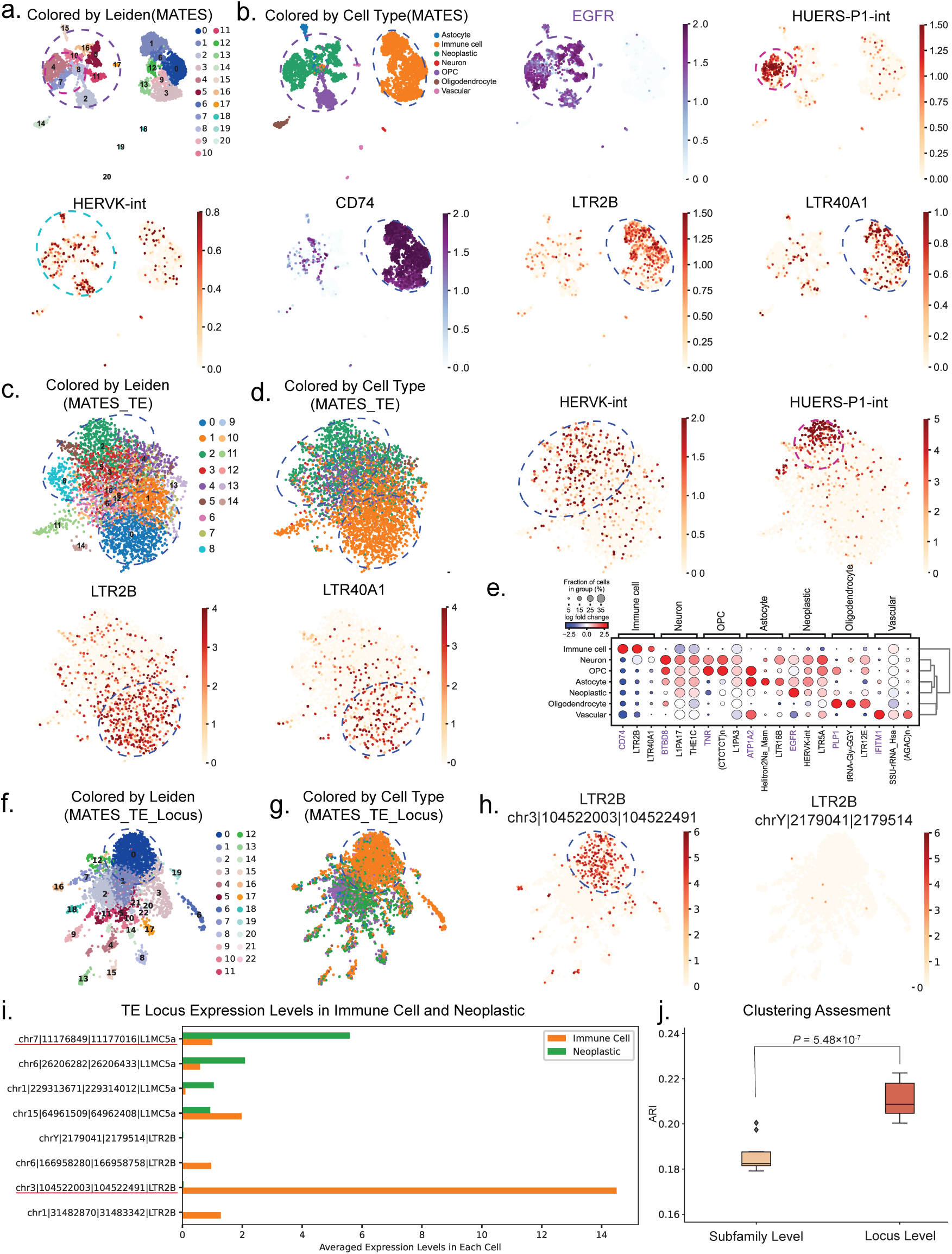
MATES quantifies disease-related TE expression in Smart-Seq2 single-cell RNA-seq data. (a-b) UMAP plots showcase cell clustering informed by gene and TE markers. “MATES” or “Gene+TE” signifies combined gene expression with TE data quantified by MATES. Initially, MATES UMAPs are colored by Leiden clusters (a), then by cell type, neoplastic (*EGFR*, HUERS-P1-int, and HERVK-int), and immune cell markers like *CD*74, LTR2B, and LTR40A1 (b). (c-d) UMAPs based solely on MATES-quantified TE expression are colored by Leiden clusters (c) and cell type with specific markers (HERVK-int, etc.) (d). (e) Dot plots elucidate the correlation between marker genes, TEs, and cell types, as identified by MATES. (f-h) Illustrate the enhanced clustering accuracy using MATES’s locus-level TE quantification, with (f) detailing Leiden clusters and (g) showing cell types. (h) The plot lists a highly expressed TE marker (LTR2B) for immune cells at the locus level and the same TE’s non-expressed locus, demonstrating MATES’s capability in locus-level TE quantification. (i) A bar plot visualizes the average locus-specific TE expression levels in immune and neoplastic cells. (j) A box plot comparison of cell clustering efficacy,Adjusted Rand Index (ARI), between MATES’s locus-level and subfamily-level TE quantification reveals the method’s enhanced resolution, demonstrating its performance in biomarker identification and cellular classification. The experiment runs with N=10 different seeds and the p-value was calculated using a one-sided Student’s t-test.

Further demonstrating MATES’s precision, we also performed cell clustering solely based on the TE count matrix quantified by MATES. While the TE-only analysis may not achieve better clustering accuracy compared to the combined analysis, it is crucial to emphasize that TE quantification holds biological information capable of producing coherent results with conventional gene-based analysis. Specifically, we systematically compared the TE-only results with gene expression clustering results and found notable similarity. Leiden clusters 0 and 1 correspond to immune cells, while clusters 2, 3, and 4 correspond to neoplastic cells (Supplementary Fig.S2 d,f). The ARI (median is 0.105, *P* = 1.03 × 10*^−^*^2^) and NMI (median is 0.161, *P* = 7.60 × 10*^−^*^4^) scores indicate a weak yet significant agreement between TE expression clustering and gene expression clustering (Supplementary Fig.S2 e). The confusion matrix further compares TE clusters to gene clusters and cell types, showing that TE cluster 0 overlaps significantly with gene clusters 0 and 1, which are primarily composed of immune cells, while TE cluster 2 aligns with gene clusters 4 and 5, containing mostly neoplastic cells (Supplementary Fig.S2 f). This suggests that TE-based clustering can accurately recapture all major cell populations, identifying their associated TE markers(Fig.3 c,d). The dot plots (Fig.3 e) not only displayed the associations between specific marker genes, TEs, and cell types but also quantified their relative expression levels, adding a deeper dimension to the data analysis.

In addition to analyzing TE expressions at the subfamily level shown above, MATES’ locus-level TE quantification offered a more comprehensive view of the cellular landscape (Fig.3 f-h, Supplementary Fig.S3). This approach facilitated the identification of highly expressed TE loci corresponding to marker TEs previously identified at the subfamily level. Notably, even for the same TE subfamily such as LTR2B, distinct loci could exhibit different expression patterns (Fig.3 h,i), emphasizing the critical necessity for precise and locus-specific TE quantification. The LTR2B locus at *chr*3|104522003|104522491|*LTR*2*B* (chrom|start|end|TE), a highly expressed TE loucs marker for immune cells, is close to the *CD*166 gene, suggesting potential regulatory interactions. *CD*166, crucial for immune cell adhesion and function [41], may be influenced by LTR2B through its regulatory elements. TEs can impact nearby gene expression by providing promoters, enhancers, and transcription factor binding sites, facilitating rapid and dynamic gene expression changes vital for immune responses [42]. Additionally, TEs are targets for epigenetic modifications, further regulating nearby genes and enhancing immune cell adaptability [43]. Further experimental analysis is needed to fully understand their interactions. In addition, the application of this locus-specific TE quantification significantly improved cell clustering accuracy compared to its subfamily-level counterpart, as demonstrated in Fig.3 j (*P* = 5.48 × 10*^−^*^7^), highlighting its critical role in analyzing cellular heterogeneity and understanding TE functions and its superiority over conventional subfamily level analysis. Please refer to the supplementary data table 1 for the identified top TE locus markers and their nearby interacting genes for neoplastic and immune cells.

Our results affirm the robustness of MATES when applied to full-length scRNA-seq data, emphasizing its effectiveness for in-depth cellular analysis within these single-cell RNA-seq datasets across varying sequencing platforms. While some existing methods (e.g., scTE) can be adapted to handle full-length single-cell RNA-seq data, their performance is often suboptimal, highlighting the value of MATES’ ability to process and interpret these datasets effectively (Supplementary Fig.S4).

### Applicability of MATES across species

MATES can quantify TE expressions not only in mammals, such as humans and mice as demonstrated above, but also in non-mammalian species, showcasing its cross-species applicability. To comprehensively evaluate MATES’ generalizability, we applied it to single-cell RNA-seq datasets from the non-mammalian species Arabidopsis thaliana (Arabidopsis) [44] and Drosophila melanogaster (Drosophila) [45]. Here we used conventional gene expression-based cell embedding and clustering as the baseline to evaluate the quality of TE quantification of different tools on these non-mammalian species.

Supplementary Fig.S5 illustrates cell clustering based on TE and gene expression in Arabidopsis using MATES, scTE, and SoloTE. The Supplementary Fig.S5 a shows the UMAP visualization of TE expression quantified by scTE compared with gene expression clusters, with an ARI of 0.3315 (*P <* 1 × 10*^−^*^6^). Panel (b) presents the UMAP visualization of TE expression quantified by SoloTE, with an ARI of 0.3200 (*P <* 1 × 10*^−^*^6^). Panel (c) shows the UMAP visualization of TE expression quantified by MATES compared with gene expression clusters, achieving an ARI of 0.3514 (*P <* 1 × 10*^−^*^6^). When combining the gene expression and TE expression quantified by computational methods (Supplementary Fig.S5 a-c), MATES achieved a 0.6668 ARI score (*P <* 1 × 10*^−^*^6^), which is higher than SoloTE (ARI = 0.5916, *P <* 1 × 10*^−^*^6^) and scTE (ARI = 0.5775, *P <* 1 × 10*^−^*^6^). This is categorized as moderate similarity, as demonstrated in a comprehensive single-cell RNA-seq clustering comparison evaluation study, where varying levels of differential expression (DE) were employed to assess clustering similarity [46]. The moderate similarity between the UMAP visualizations in panels (a), (b), and (c) indicates that TE-based clustering mirrors gene expression-based clustering, with MATES showing the highest agreement. Here, we employed permutation tests to calculate p-values associated with the similarity ARIs calculated above (see Methods for details). Additionally, the identification of marker TEs using MATES further supports its effectiveness. Panel (d) highlights that specific marker TEs were identified for various cell populations. For instance, marker gene *KCS*6 identified the outer cell layer, while *HIK* identified partially dividing and inner cell layers. Correspondingly, marker TEs ATCopia66LTR and ATCopia41LTR were identified for the outer cell layer and the partially dividing and inner cell layers, respectively.

For Drosophila, Supplementary Fig.S6 shows clustering results based on TE and gene expression using MATES and scTE. UMAP visualizations show TE expression quantified by MATES (ARI = 0.3188, *P* = 4.60 × 10*^−^*^5^, Supplementary Fig.S6 a) and by scTE (ARI = 0.3078, *P* = 4.60 × 10*^−^*^5^, Supplementary Fig.S6 b). This is categorized as moderate similarity, as supported by studies assessing clustering similarity through varying levels of DE [46]. The moderate similarity between the UMAP visualizations in panels Supplementary Fig.S6 a,b indicate that TE-based clustering mirrors gene expression-based clustering, with MATES showing slightly higher agreement. Additionally, the identification of marker TEs using MATES further supports its effectiveness. Supplementary Fig.S6 c highlights that specific marker TEs were identified for various cell populations. For example, the marker TE GYPSY12 LTR was identified for cluster type 8, as defined by conventional gene clustering.

These findings, based on both the moderate ARI values indicating significant clustering similarity, as evidenced by the permutation test p-values, and the effective identification of marker TEs for specific cell types, demonstrate MATES’s robustness and applicability across both mammalian and non-mammalian species.

### Applicability of MATES across modalities

Beyond the general applicability to various species as shown above, our proposed MATES model also demonstrates its versatility by effectively quantifying TEs in not only transcriptome data but also epigenome data. To validate this adaptability, we applied MATES to a 10x single-cell ATAC-seq dataset of adult mouse brain [47]. This compatibility with single-cell data across different modalities is crucial, as many existing methods are tailored exclusively for transcriptomic data and may not perform as effectively with other single-cell modalities. MATES enables the quantification of TE locus-specific attributes, such as chromatin accessibility, across diverse single-cell data modalities, extending its utility beyond transcriptomics.

By quantifying chromatin accessibility at the TE subfamily level and incorporating standard single-cell ATAC-seq peaks, MATES facilitates refined cell clustering and the identification of TE markers that are characteristic of distinct cell populations (Fig.4 a,b, Supplementary Fig.S7 a). TEs such as RMER16_MM and RLTR44B from the ERVK family show exclusive accessibility in macrophages [48, 49], while MamRep434 and MER124 are preferentially accessible in astrocytes [50], underscoring TEs’ significant roles in neurogenesis and astrogliogenesis [51]. For example, MamRep434’s significant contribution to the motif of Lhx2, a key transcription factor in neurogenesis, is emblematic of the functional implications of TE accessibility in cell identity and function [50, 52]. Leveraging solely the TE expression data quantified by MATES for clustering, we successfully identified TE biomarkers and maintained clear delineation among cell groups (Fig.4 c,d, Supplementary Fig.S7 b), confirming that the insights gleaned from TE quantification are rooted in genuine biological phenomena. Although integrating TE expression with conventional quantification like gene or peak counts generally produces the best cell embedding quality and clustering, TE-only analysis is essential for highlighting the specific contributions of TEs beyond conventional gene or peak expressions. By comparing TE-based results with conventional gene or peak-based results, we demonstrated that TE quantification is highly informative and can produce consistent cell clustering results similar to conventional analyses (Supplementary Fig.S2 g-i). The median ARI between TE-based clusters and gene-based clusters is 0.309 (*P* = 5.60 × 10*^−^*^5^), and median NMI is 0.438 (*P* = 4.60 × 10*^−^*^6^).

**Fig. 4.**
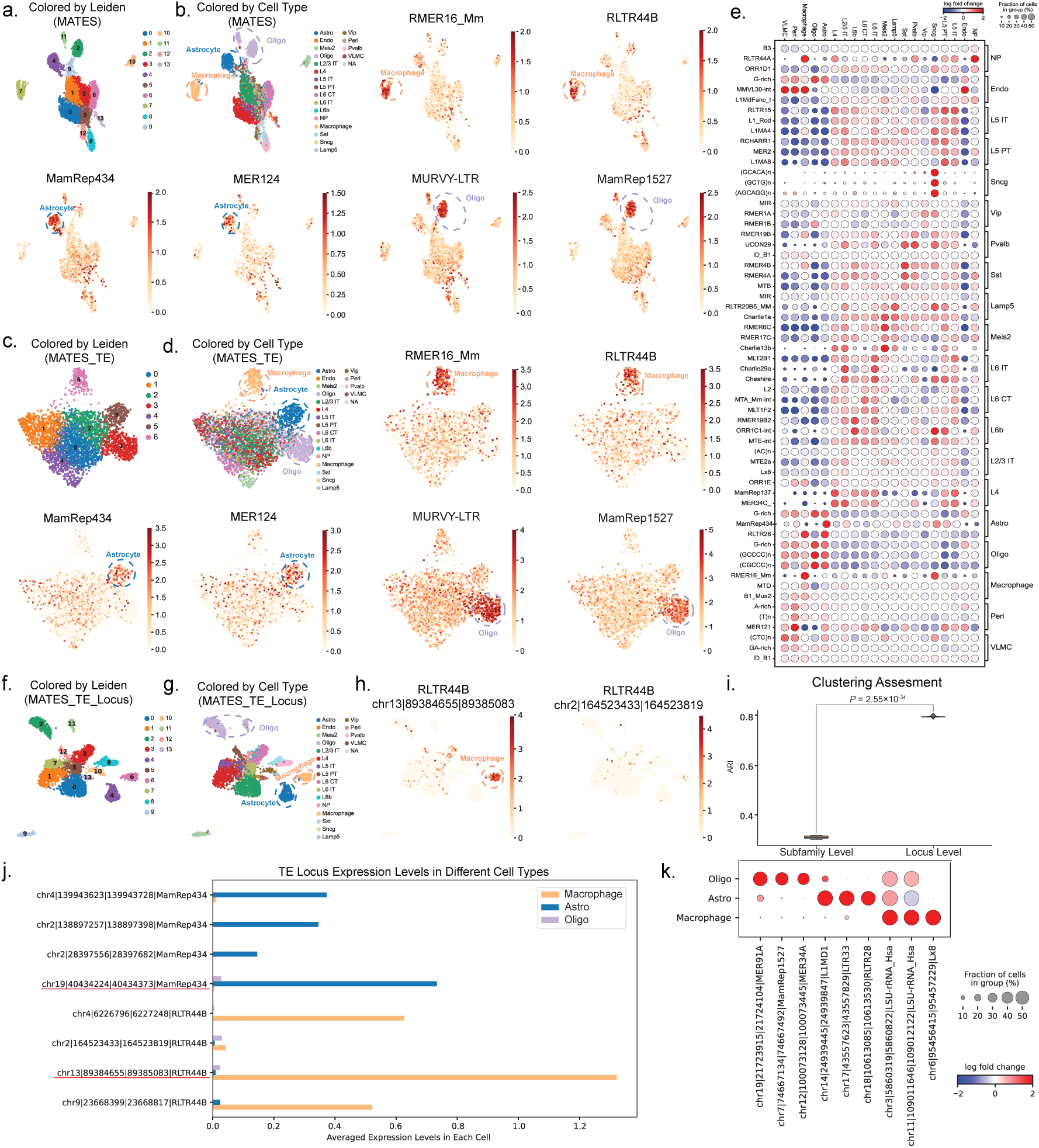
MATES versatility application on adult mouse brain scATAC-seq data. (a-d) UMAP plots demonstrate the effectiveness of MATES quantification in cell clustering and identifying signature TE markers, using both TEs and Peaks for clustering, with (a) displaying Leiden clusters, and (b) illustrating cell types and TE markers. Key TE biomarkers such as RMER16_Mm, RLTR44B in macrophages, MamRep434, MER124 in Astrocytes, and MURVY-LTR, MamRep1527 in oligodendrocytes are identified. (c-d) Showcase the specificity of MATES in TE-centric clustering (using only TE quantification by MATES), with (c) focused on Leiden clusters, (d) on cell types and the distinctive TE markers previously noted. (e) Dot plot concisely presents cell type-specific TE markers uncovered by MATES. (f-h) These panels illustrate MATES’s improved clustering accuracy using the locus level TE quantification. Panel (f) features UMAP visualizations based on locus-level TE quantification, with colors representing Leiden clusters. Panel (g) displays the same UMAP, but with color coding to differentiate various cell types. Panel (h) provides a specific example of the TE marker RLTR44B in Macrophages at the locus level, in contrast to a non-open locus of the same TE, demonstrating MATES’s capability in detailed locus-level TE quantification. (i) The box plot in panel i contrasts the cell clustering efficacy, Adjusted Rand Index (ARI), of MATES when comparing locus-level versus subfamily-level TE quantification. This comparison highlights the advantages of employing locus-specific TE quantification. The experiments run with N=10 different seeds and the associated p-value was determined using a one-sided Student’s t-test. (j) The bar plot visualizes the average locus-specific TE expression in Macrophage, Oligo, and Astro cells. (k) The dot plot shows the locus-specific TE markers, identified by MATES, for individual cell types. types, as identified by MATES.

The identified signature TEs for all cell populations were shown in Fig.4 e. Beyond subfamily-level TE quantification and analysis, MATES also delivers locus-specific TE quantification, identifying signature TEs for each cell population with precise TE locus positions (Fig.4 f-h, Supplementary Fig.S7 c,d). Locus-level TE quantification demonstrates significantly higher (*P* = 2.55 × 10*^−^*^34^) cell clustering accuracy compared to subfamily-level analysis (Fig.4 i). This underscores the effectiveness of MATES in locus-level TE quantifications and their potential benefits for understanding cellular states within the data. Marker TEs identified at the locus level align with those detected at the subfamily level, validating the method’s accuracy and yielding locus-specific insights into TE’s impact on chromatin accessibility [53]. Supplementary Fig.S7 provides additional supporting evidence of MATES’ effectiveness in the 10x scATAC-seq dataset, further solidifying its position as a versatile tool for TE quantification and analysis at the single-cell level across different modalities.

These locus-level TE biomarkers for each cell population can potentially reveal interactions with nearby genes and their impact on regulating cellular states for specific cell types (Fig.4 j,k). For example, in this mouse brain scATAC dataset, the locus chr13|89384655|89385083|RLTR44B exhibits substantial chromatin accessibility in cells that should be annotated as microglia, despite their previous macrophage annotation (Fig.4 h). One of the flanking genes is *Edil3*. Quantitative proteomics indicate increased *Edil*3 expression in isolated microglia of APP-KI mice compared to WT mice [54]. This locus is also located upstream of the gene *Hapln*1, which has been recently identified as a macrophage-related regulator and is correlated with cancer immunotherapy [55]. These findings support the potential roles of this RLTR44B locus in the macrophage/microglia cell population. Similarly, the locus chr19|40434224|40434373|MamRep434 (Supplementary Fig.S7 c), located near the *Sorbs*1 gene associated with astrogliosis, demonstrates elevated expression in individuals with schizophrenia who express high levels of inflammatory markers [56]. Additionally, genes situated within a 200kb flanking region of this transposable element include *Aldh*18*a*1 [57] and *Entpd*19 [58], which play pivotal roles in astrocyte functions and their interactions with non-astrocytic cells. These findings collectively underscore the regulatory potential of these identified marker TE loci in influencing astrocyte-related gene expression. Another notable case is the locus chr18|10613085|10613530|RLTR28, which shows high accessibility in scATAC-seq and could be a potential enhancer for the nearby stress-responsive gene *Abhd*3 in astrocytes, highlighting the role of TEs in stress response through genetic mutations and epigenomic variations [59]. Additionally, MATES identified the locus chr17|43557623|43557829|LTR33 near the Phospholipase A group 7 (*Pla*2*g*7) gene, suggesting it might serve as a tissue-specific expression enhancer of this gene in cortical GM astrocytes. This finding is supported by the reported predominant expression of *Pla*2*g*7 in astrocytes [60]. Please refer to the supplementary data table 2 for the top identified TE locus markers and their nearby interacting genes for Macrophages, Astro and Oligo.

Overall, these examples illustrate how MATES enables detailed interrogation of interactions between TE loci and nearby genes, potentially unlocking deeper understanding of biological mechanisms and providing valuable insights into the regulation of cellular states.

### Consistency between single-cell TE quantification by MATES and corresponding bulk quantification via conventional methods

To evaluate the robustness of MATES, we compared both scRNA-seq and scATAC-seq TE quantification with matching bulk datasets. MATES, optimized for single-cell data, demonstrated consistent results across various experimental configurations. For this comparison, bulk data were generated from 10X scRNA-seq and scATAC-seq (pseudo-bulk), simulating bulk RNA-seq and bulk ATAC-seq datasets. Utilizing pseudo-bulk data derived from the same set of cells minimized potential batch effects, ensuring a rigorous comparison. This approach facilitated a direct comparison between single-cell TE expression quantified by MATES and bulk-level quantification from pseudo-bulk data using TEtranscripts [11] and Telescope [12], which are specifically designed for bulk TE quantification. Please refer to ‘MATES comparison with existing TE quantification methods on bulk data’ in the Methods section for our strategy to compare the single-cell TE quantification from MATES with the corresponding bulk TE quantification from existing approaches.

The results for RNA data, illustrated in Supplementary Fig.S8, demonstrate a high degree of correlation between pseudo-bulk TE expression quantified by MATES and the ground truth. Specifically, MATES showed a strong correlation with TEtranscripts (*R*^2^=0.9429; Supplementary Fig.S8 a) and with Telescope (*R*^2^=0.9664; Supplementary Fig.S8 b), indicating robust performance. To provide a comprehensive comparison regarding TEs and repetitive elements, we included only well-defined TEs (DNA, LTR, RC, Retroposon, SINE, and LINE) in the analysis of pseudo-bulk RNA data.

The results for comparisons made on ATAC data, depicted in correlation scatter plots (Supplementary Fig.S9), also indicate a high degree of correlation between MATES and TEtranscripts (*R*^2^ = 0.9901; Supplementary Fig.S9 a) as well as between MATES and Telescope (*R*^2^ = 0.9890; Supplementary Fig.S9 b). These high correlation coefficients affirm that MATES reliably captures TE expression profiles comparable to those obtained from bulk ATAC-seq data. Additionally, selective genome views (Supplementary Fig.S9 c) highlight specific TE regions, further demonstrating the concordance between single-cell (MATES) and bulk (TEtranscripts, Telescope) TE quantifications. This analysis incorporated repetitive elements to ensure comprehensive evaluation.

These results underscore the efficacy of MATES in providing consistent and reliable TE quantification at the single-cell level, on par with conventional bulk methods. This consistency not only validates the accuracy of MATES but also enhances our understanding of TE dynamics at the single-cell level, potentially unlocking deeper insights into underlying biological mechanisms.

### MATES enables multi-omic TE quantification and analysis

To further attest to the broad applicability of MATES, we applied it to a single-cell multi-omics dataset (10x multiome) [61]. By amalgamating the TE quantification provided by MATES with conventional gene expression data (RNA transcripts from scRNA-seq) and matched accessibility quantification (chromatin accessibility from scATAC-seq), we distinguished various cell populations and their associated markers, as depicted in Fig.5 a,b.

**Fig. 5.**
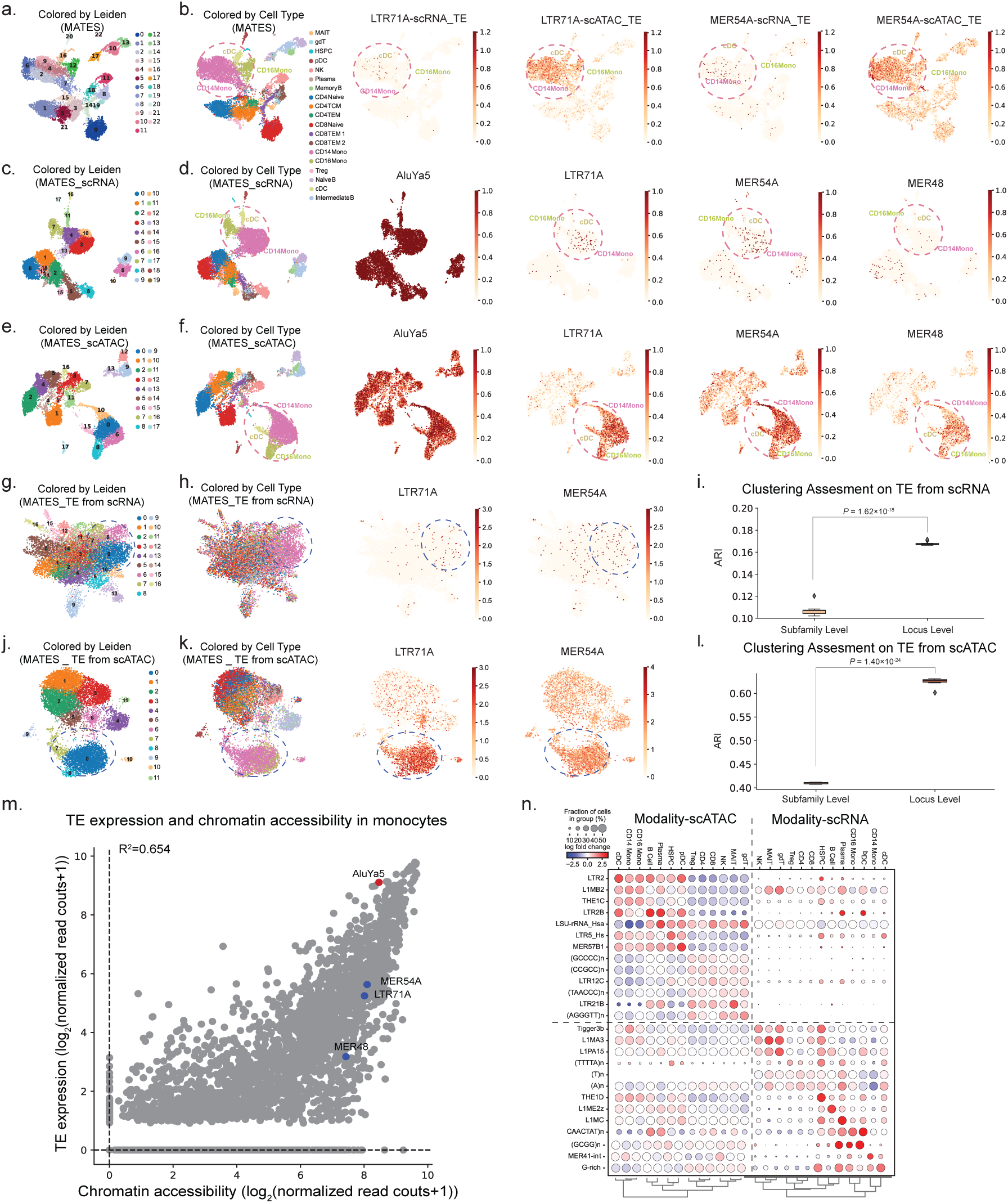
Multi-modal TE analysis in human PBMCs using MATES. (a-b) UMAP plots of joint MATES clustering using both sc_RNA and sc_ATAC modalities, with (a) illustrating Leiden clusters and (b) detailing cell type clusters. (c-f) TE quantification across modalities highlights the complementarity of multi-modal TE quantification. Here, (c-d) feature UMAP clustering by Gene and TE in the scRNA modality, while (e-f) display UMAP clustering by Peaks and TE in the scATAC modality. Both (c) and (e) are colored by Leiden clusters, and (d) and (f) by cell types, showcasing the differential expression of TEs like AluYa5 across both modalities, whereas MER48, LTR71A, and MER54A appear specific to scATAC. (g-l) This series of UMAP plots and box plots illustrates the multi-modal TE analysis. (g) and (j) feature UMAP plots of TE expression, colored by Leiden clusters to highlight the clustering patterns. (h) and (k) are UMAP plots that focus on different cell types and TE markers, providing insights into the distinct identities of cells and their associated TEs. (i) and (l) are box plots contrasting the cell clustering effectiveness, Adjusted Rand Index (ARI). The box plots underscore the improved resolution provided by locus-level quantification over subfamily-level quantification. The experiments in these two plots run with N=10 different seeds and the p-values were calculated using a one-sided Student’s t-test. (m) Illustrates TE biomarkers discerned via scRNA and scATAC modalities, noting that high expression TEs often correlate with increased chromatin accessibility, while the reverse is not uniformly observed, highlighting the unique contributions of each modality. (n) A dot plot captures signature TEs per cell type, validating the complementary nature of scATAC and scRNA data for a holistic view of transposon dynamics.

MATES’s ability to cluster diverse cells, identifying distinct cell populations and TE biomarkers across different sequencing techniques, introduced MATES’ potential in harnessing multi-omics data for in-depth cell analysis (Fig.5 c-f), the results underscore the synergistic interplay of different modalities within MATES. Notably, when solely relying on MATES’ TE quantification (TE scRNA and TE scATAC), the method effectively captured the primary cell populations and their signature TE markers. In contrast, quantification at the locus level further improved clustering performance(Fig.5 g-l, Supplementary Fig.S10 a-d). Analyzing the multi-omics data, MATES unexpectedly reveals that certain TEs are uniquely discernible in specific cell sub-populations and exclusive to a particular modality (Fig.5 m,n, Supplementary Fig.S10 e). TEs with elevated gene expression often exhibit increased chromatin accessibility (represented by red dots, such as AluYa5, have been reported [62]). Conversely, transposons with enhanced chromatin accessibility do not always indicate high expression levels (marked as blue dots). For instance, LTR71A is predominantly detected via scATAC-seq, while not present in the scRNA-seq dataset. A similar trend is observed with TE biomarkers like MER48, and MER54A, suggesting these TEs are accessible in uninfected monocytes but transcriptionally dormant. This leads us to propose that such TEs may be primed to a “poised” state that might correlate to monocyte function [63]. Notably, several of these TEs, previously identified as “poised”, featured prominently among the upregulated TEs. This enrichment calculated by hypergeometric test, with a significant p-value of *P* = 5.91 × 10*^−^*^24^, was determined by comparing the marker TEs identified through MATES with those upregulated TEs reported in the study (Supplementary Fig.S10 f).

This observation highlights the capability of MATES to uncover biological features of TE. Here, our result emphasizes that when analyzed through the prism of TEs, chromatin accessibility, and RNA abundance offer complementary insights into the single-cell state (see Fig.5 m). The strength of MATES is demonstrated in Supplementary Fig.S10 g. Within this framework, TE quantification by MATES boosts cell clustering accuracy and helps to build new hypotheses.

### Method benchmarking in single-cell TE quantification

In our study, we conduct a detailed benchmarking of the MATES approach for TE quantification and its effect on cell clustering in single-cell datasets with various modalities. This is compared with the two established methods, scTE and SoloTE. Our benchmarking primarily concentrates on TE quantification at the subfamily level, due to the limitations of existing methodologies. Both scTE and SoloTE are incapable of providing locus-specific TE quantification that accounts for multimapping reads. Specifically, scTE is confined to subfamily level quantification, while SoloTE does offer locus-specific quantification but is constrained to unique mapping reads. The prevalence of multimapping reads among TEs highlights the critical need for accurate locus-specific TE quantification to enhance our understanding of cellular states. To ensure a fair comparison in our benchmarking, we have restricted our analysis in this section to the subfamily level, a feature shared by both scTE and SoloTE. However, the advantages of MATES in providing more accurate locus-specific TE quantification over the subfamily level have been discussed in earlier sections of our study for each individual dataset.

Here, we perform comprehensive benchmarking of TE quantification methods, evaluating them based on their downstream cell clustering performance. We analyze TE information from three distinct perspectives to illuminate the various aspects of TE expression’s relevance and utility. Firstly, we analyzed gene/peak and TE expression concurrently to demonstrate how TE expression complements conventional gene/peak quantifications from single-cell data. This analysis underscores the added value that TE data can provide alongside traditional metrics. Secondly, we focused exclusively on TE expression to showcase the inherent power and information encapsulated within TE data alone. This approach aimed to illustrate that TE quantification, independent of conventional gene/peak data, holds sufficient information for effective cell clustering. Lastly, we assessed TE expression based on multi-mapping reads to emphasize the ability of the MATES method in handling these reads for TE quantification. This aspect was particularly crucial for understanding the robustness of MATES in dealing with the complexities of TE data. These comparisons were referred to as Gene/Peak + TE expression (Gene+TE/Peak+TE), TE expression (TE), and Multi-mapping TE expression (Multi TE), respectively.

The benchmarking in this work included single-cell RNA datasets obtained using both the 10x and Smart-Seq2 platforms, as well as single-cell ATAC sequencing (scATAC) datasets. It is important to note that SoloTE is not compatible with Smart-Seq2 and scATAC single-cell data. Consequently, SoloTE was excluded from the comparative analysis for these two datasets. To assess the clustering results derived from the three perspectives outlined earlier, ARI and NMI as our key benchmarking standards. Our analysis compared MATES with two existing single-cell TE quantification methods scTE and SoloTE, which primarily differ in their approach to handling multi-mapping reads. We observed significant differences in their performance: scTE uses a basic correction method, SoloTE selects the top-scoring alignment, and MATES utilizes a probabilistic approach informed by local read context. The statistical significance of our results, indicated by p-values, reinforces the improved performance of MATES in TE quantification, as evidenced by improved cell clustering performance compared to existing methods.

In the analysis that integrated both Gene and TE expression, MATES demonstrated better performance than scTE and SoloTE, especially evident in the 10x scRNA dataset for chemical reprogramming. In this context, scTE’s effectiveness was markedly inferior compared to a gene expression-only approach, highlighting the importance of accurate multi-mapping read assignment. This was evidenced by MATES achieving higher ARI and NMI scores across all datasets tested, as shown in the left panels of Fig.6 a,b. To further showcase and compare the accuracy of TE quantification by different methods, we conducted a cell clustering analysis based on only the TE quantification, excluding gene expression data. Here again, MATES outshone scTE and SoloTE in terms of clustering efficiency, which was reflected in improved ARI and NMI scores (see the middle panels of Fig.6 a,b). This result underscores the capability of MATES in handling multi-mapping reads and demonstrates its significance in TE quantification. In scenarios focusing exclusively on multi-mapping TE reads, MATES manages these challenging read assignments became apparent. MATES consistently outperformed scTE and SoloTE in these tests, as depicted in the middle and right panels of Fig.6 a,b. This consistent improvement across different testing conditions demonstrates the robustness and effectiveness of MATES in TE quantification. Its strength in the assignment of multi-mapping reads contributes to the accuracy in subsequent cell clustering tasks.

**Fig. 6.**
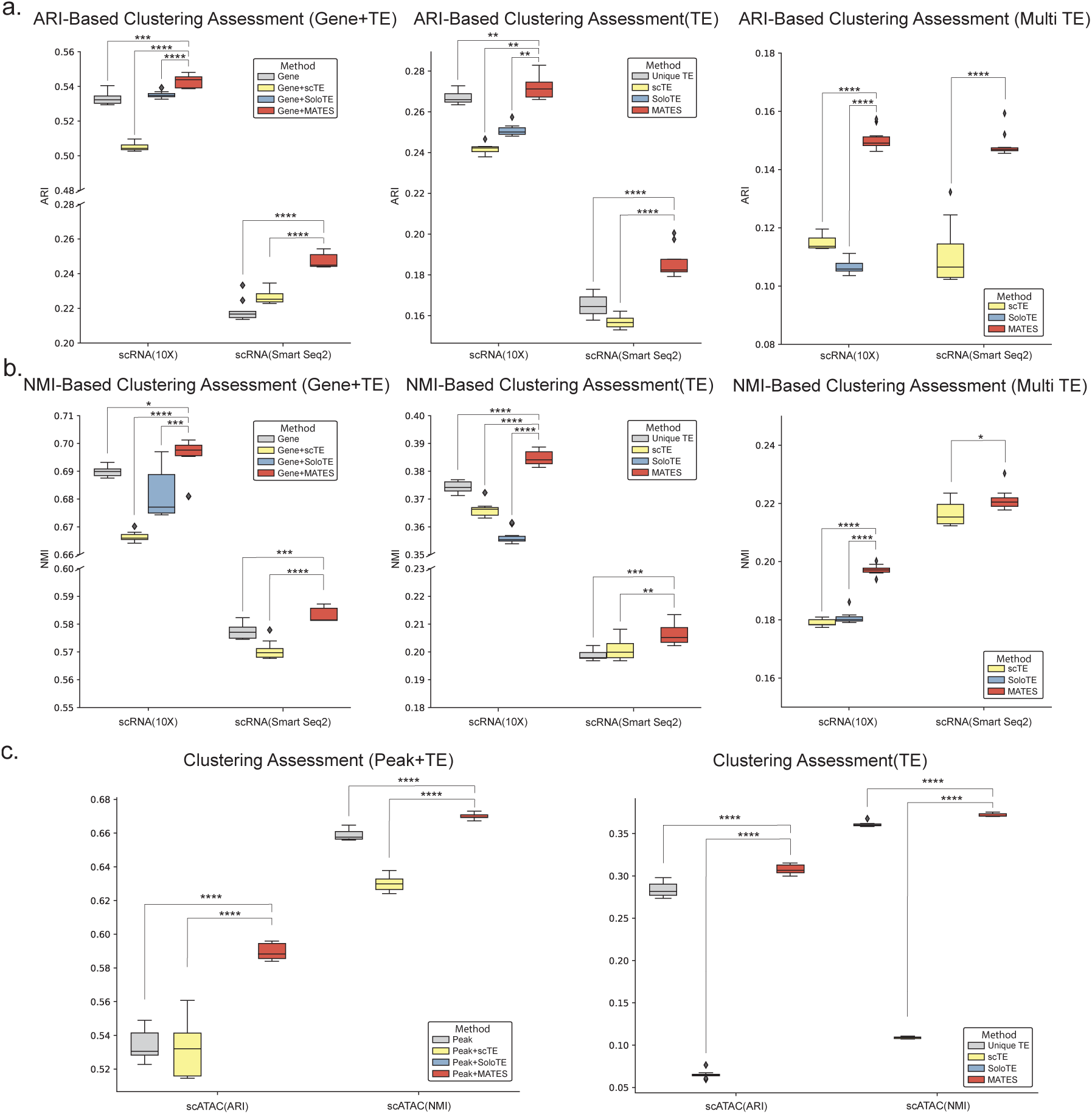
Benchmarking MATES performance in diverse single-cell datasets. This assessment highlights MATES’s efficiency in cell clustering, evaluated through the Adjusted Rand Index (ARI) and Normalized Mutual Information (NMI), using different TE quantification strategies. (a-b) The impact of various TE quantification methods on cell clustering is compared within the 10x scRNA Dataset of Chemical Reprogramming and Smart-Seq2 Dataset of Glioblastoma. These methods include scTE (gene expression combined with scTE-quantified TE), SoloTE (gene expression combined with SoloTE-quantified TE), and MATES (gene expression integrated with MATES-quantified TE). MATES outperforms both scTE and SoloTE by enhancing gene expression with TE data (left). The middle panel compares clustering based solely on TE quantification methods—unique TEs, scTE, SoloTE, and MATES—with ’unique TEs’ representing unique-mapping reads TE expression, highlighting MATES’s consistently improved performance. The right panel confirms MATES’s advantage over scTE and SoloTE when considering only multi-mapping TE reads. Note: SoloTE’s incompatibility with Smart-Seq2 data results in a blank section. Panel (a) uses ARI for evaluation, while panel (b) utilizes NMI. (c) The 10x scATAC Dataset of the Adult Mouse Brain is analyzed to contrast peak and TE quantification using scTE and MATES against peak-only datasets (left). TE mapping reads from scTE and MATES are also compared against unique TE mapping reads (right). SoloTE’s incompatibility with scATAC data leads to its exclusion from this part of the analysis. (The experiments run with N=10 different seeds. The p-values were calculated using a one-sided Student’s t-test, with significance levels indicated as * for *P <* 0.05, ** for *P <* 0.01, *** for *P <* 0.001, and **** for *P <* 0.0001).

In our benchmarking analysis of the 10x scATAC mouse brain dataset, we assessed the MATES approach against scTE and peak-only quantification methods. This evaluation aimed to determine the cell clustering accuracy when combining peak and TE data. Our results indicated a clear advantage of MATES over both scTE and peak-only methods in terms of clustering accuracy, showcasing its capacity to integrate additional chromatin accessibility insights effectively. A key aspect of MATES’s performance was its comprehensive quantification of TE expression, which encompassed both unique mapping reads and overall TE data. This approach significantly surpassed scTE, contributing to an enhanced clustering process. The inclusion of multi-mapping reads by MATES, although less critical than in scRNA datasets due to its lower frequency in this scATAC-seq data, provided valuable insights to the clustering analysis, as illustrated in Fig.6 c. These findings not only highlight the effectiveness of MATES but also demonstrate its adaptability and potential for broad applications in various single-cell data modalities.

Furthermore, we benchmarked MATES against existing methods for locus-level TE quantification using the 2CLCs 10X scRNA dataset. Since scTE does not support locus-level TE quantification, we only compared the results from MATES with those from SoloTE. Supplementary Fig.S11 shows the ARI scores for cell clustering based on locus-level TEs, where MATES achieved a 10.52% higher ARI score than SoloTE (*P* = 2.60 × 10*^−^*^12^). This improved performance may be attributed to MATES’s ability to leverage locus-level TE expression from multi-mapping reads, whereas SoloTE can only quantify locus-level TE expression from uniquely-mapped reads.

### Validation of MATES quantification accuracy with single-cell long-read sequencing data and simulation experiments

#### Validation through long-read sequencing data

To validate the accuracy of our TE quantification method, MATES, we used long-read sequencing data from both PacBio and Nanopore platforms. PacBio’s Sequel II system generates high-fidelity (HiFi) reads with superior accuracy, often exceeding 99%, making it ideal for applications requiring precise base-level resolution, such as identifying specific TE insertion sites [64]. Nanopore sequencing, although typically known for its ultra-long reads, provided reads around 900 bp in this study. With an accuracy rate ranging from 85% to 95%, it still offers valuable insights into repetitive regions and TE structures [65]. Both long-read sequencing technologies enhance our ability to distinguish similar TE instances by spanning long repetitive regions and capturing more mutations. Including datasets from both platforms robustly validates MATES by leveraging the complementary strengths of each technology.

To ascertain the precision of TE quantification by MATES, we first utilized a melanoma brain metastasis dataset from nanopore sequencing platform, as detailed in Shiau et al. [66]. This dataset comprised single-cell nanopore RNA sequencing (scNanoRNAseq) [67], producing long-read data with an average length of 937 base pairs (bp). This is significantly longer than the corresponding short-read data of 222 bp obtained from 10x scRNA-seq (real sequencing reads), which came from the same set of cells in the study. The longer read length of nanopore sequencing is advantageous for TE quantification, as it allows for more accurate read alignment [68] (i.e., reduced multi-mapping rate ∼ 1% in nanopore sequencing compared to over 12% in 10x shortread sequencing from the same study), thereby enhancing the reliability of TE quantification. This level of precision from longread data serves as the ground truth for our validation of short-read TE quantification by MATES. In our validation of MATES through nanopore long-read sequencing data, we compared TE expression quantified by each method, including MATES, scTE, and SoloTE, within individual cells. To enhance the accuracy of our correlation calculations and accurately reflect the relationship between short-read and long-read expressions, we included only well defined TE families (DNA, LTR, RC, Retroposon, SINE, and LINE). This was to ensure a focused and accurate analysis of transposable elements and the *R*^2^ values truly represented the biological data, free from distortions caused by non-expressive or anomalous readings. MATES’s quantification of TEs demonstrated a strong correlation with the ground-truth long-read data (*R*^2^ = 0.7531), surpassing both scTE (*R*^2^ = 0.6499) and SoloTE (*R*^2^ = 0.6841) in quantifying TE expression at the subfamily level from the real 10x short-read data, as seen in Fig.7 a-c. Additionally, the pseudo-bulk TE expression in the data, determined by averaging the TE expression across all cells in the real short-read 10x sequencing dataset, is also quantified by each respective method. MATES demonstrated a stronger correlation with the established ground-truth long-read data, achieving an *R*^2^ value of 0.9276. This performance exceeded that of other methods such as scTE and SoloTE, which achieved *R*^2^ values of 0.8591 and 0.8654, respectively (Supplementary Fig.S12).

**Fig 7.**
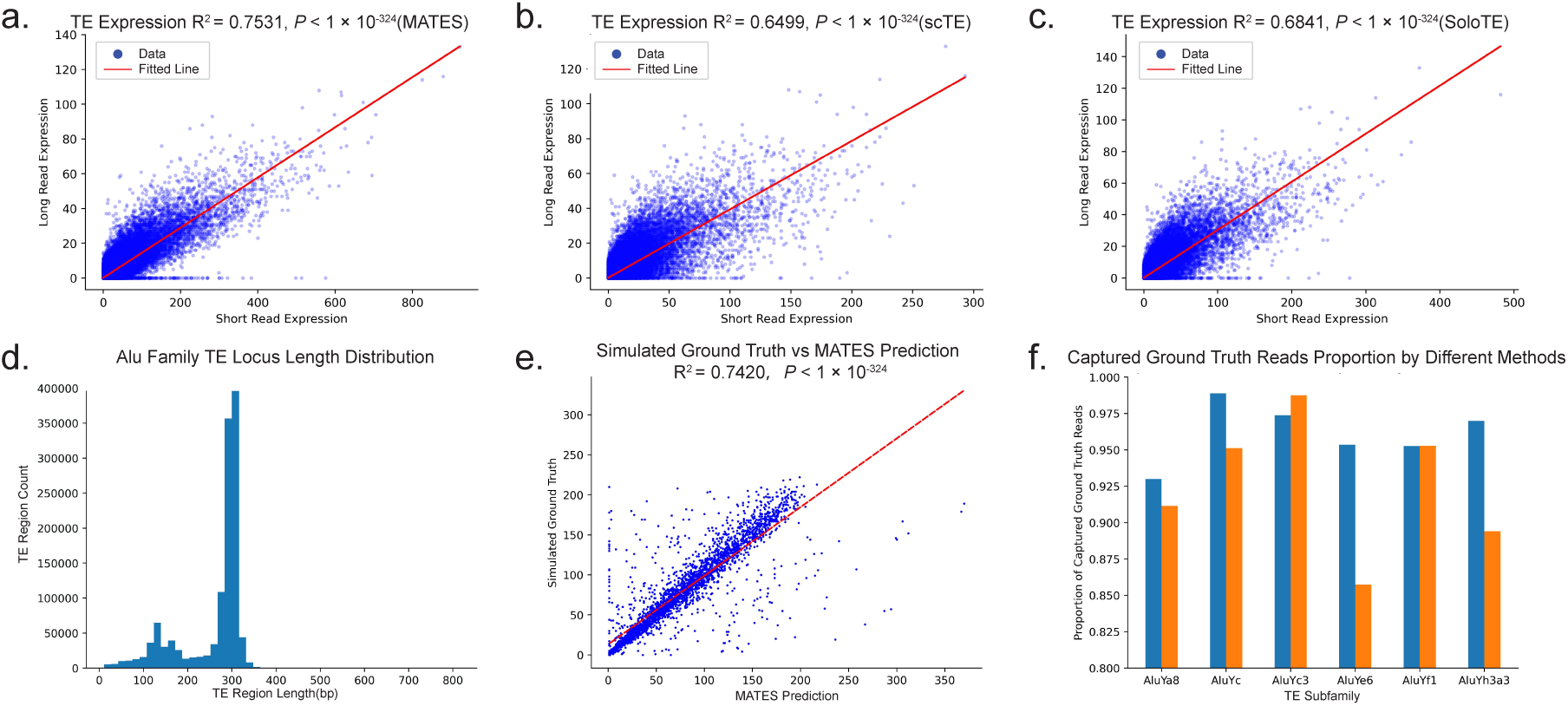
Validation of MATES TE quantification using nanopore long-read single-cell sequencing and simulation. (a-c) Analyze the correlation between TE expression quantified by (a)MATES, (b)scTE, and (c) SoloTE from real 10x short-read data and the nanopore long-read sequencing for the same set of cells. This comparison is at the subfamily level. (d) The distribution of lengths of Alu family TE regions. (e) Correlation between locus-level TE expression quantified by MATES and the simulated ground truth for the Alu repeat simulated data. (f) Comparison of quantified results by MATES and scTE to the simulated ground truth for the Alu repeat simulated data. The blue bar and orange bar represent the percentage of simulated reads captured by MATES and scTE, respectively. Among the 6 simulated Alu families, MATES on average captured 96.14% simulated reads while scTE recaptured 92.57%. The p-values of *R*^2^ (Coefficient of determination) were calculated using the one-sided F-test.

We then benchmarked MATES against scTE and SoloTE using PacBio data from the postnatal mouse brain, with the processed long-read data of 7,896 cell barcodes and an average length of 1,164 bp [69]. The data also contains short-reads data from the same number of cells with a length of 91 bp. The results from this analysis showed strong correlations between the long-read and short-read expression data, with MATES achieving an *R*^2^ value of 0.5096, compared to 0.3537 for scTE and 0.4199 for SoloTE (Supplementary Fig.S13). These findings indicate that PacBio long-read validation supports that MATES delivers a much better accuracy in TE quantification.

#### Validation through controlled simulation

Despite the validation provided by long-read sequencing from Nanopore and PacBio platforms, these technologies have limitations in capturing short TEs like Alu repeats. To address these limitations, we conducted benchmarking using simulated Alu repeats data, with length around 300 bp (Fig.7 d). Using the simulation data as the ground truth allows for more robust comparisons of different methods’ performances, as the benchmarking is not affected by sequencing errors or other technical limitations. Please see ‘Validation with controlled simulation’ in Methods for the details of constructing the simulation dataset. The quantification of MATES were then compared against the simulated ground truth, yielding an *R*^2^ value of 0.7420 (Fig.7 e). The simulated data consists of full-length RNA-seq reads without UMIs, which are required by SoloTE for processing and mapping. Consequently, SoloTE is incompatible with this data, prompting a focus on comparing subfamily-level performance with scTE. As illustrated in Fig.7 f, MATES demonstrated closer quantification to the ground truth compared to other evaluated methods.

Additionally, simple and low-complexity repeats, which are pervasive in genomic data are also short. However, simple repeats have important impacts on evolution and human disease [70–72]. Thererefore, here we also validated the effectiveness of MATES in quantifying the expression for simple and low-complexity repeats. Cluster-specific simple repeats also emerged as top marker TEs in the above results (e.g. Fig.3 e, Fig.4 e, and Fig.5 n), highlighting their potential roles. To show the accuracy of read assignment at the locus level for simple repeats, we conducted a detailed simulation study similar to the simulation of Alu family. The correlation between the MATES quantified read numbers and the simulated reads at each locus was evaluated, achieving a high correlation (*R*^2^ = 0.7479, *P* = 2.02 × 10*^−^*^43^), as illustrated in Supplementary Fig.S14 a. Supplementary Fig.S14 b compares the difference in read numbers between the ground truth and the quantified results from MATES and scTE, respectively. Shorter error bars indicate closer alignment to the ground truth. Although scTE showed better performance in a few TE instances, MATES generally demonstrated improved performance across most subfamilies.

Another concern is that repeats can form different isoforms through alternative splicing, further challenging model performance. For instance, HERV-K has many isoforms with varying lengths (Supplementary Fig.S15 a,b). To address this, we employed a controlled simulation to demonstrate MATES’ capacity for quantifying TE isoforms. Specifically, we simulated TE1 and TE2 (Supplementary Fig.S15 c) and obtained 4126 ground-truth reads for TE1 and 1282 reads for TE2. Using MATES, pre-trained on human scRNA-seq data, we predicted 4140 reads for TE1 and 1295 reads for TE2 (Supplementary Fig.S15 d). These results closely matched the ground truth, demonstrating MATES’ ability to accurately quantify TE isoforms and differentiate between regions with and without deletions, even with the complexity of alternative splicing.

To directly evaluate the accuracy of locus-level TE quantification, we simulated full-length short reads from a scNanoRNA-seq long-read data [66] to serve as a proxy ground truth. The advantage of nanopore long-read sequencing lies in its ability to generate reads that are sufficiently long to capture variations between different TE instances of the same or similar TE subfamilies. This capability provides a more precise representation of locus-level TE quantification, enabling a systematic and objective assessment of the performance of methodologies in quantifying TE loci. MATES and the SoloTE strategy were run on the simulation data and evaluated against the proxy ground truth from the long-read data. As a result, MATES demonstrated a higher correlation (*R*^2^ = 0.4923*, P <* 1 × 10*^−^*^324^) in profiling locus-specific TE expression compared to the imitation of SoloTE strategy (*R*^2^ = 0.1178*, P <* 1 × 10*^−^*^324^) (Supplementary Fig.S16). This result highlights MATES’s better performance in profiling locus-specific TE expression.

In conclusion, these long-read sequencing and controlled simulation based validations effectively demonstrated MATES’s robustness in quantifying TEs, even in complex scenarios involving alternative splicing and short TEs. This underscores the capability of MATES to handle diverse and challenging contexts in TE quantification, providing accurate and meaningful insights.

## Discussion

To address the multi-mapping read assignment challenge in accurate TE quantification, here we introduce MATES, a deep neural network framework tailored for quantifying TE expressions at the single-cell level, with a particular focus on multi-mapping TE reads. Utilizing an AutoEncoder, MATES learns the distribution patterns of unique mapping reads at individual TE loci based on the coverage vectors from unique-dominant TE regions. It integrates both unique-mapping and multi-mapping reads to accurately quantify TE expression at the locus level. When applied to various datasets, including 10x and Smart-Seq2 scRNA-seq data, 10x scATAC-seq data, and 10x Multiome data, it consistently outperformed existing methods. This tool is not limited to subfamily-level TE quantification but also delivers locus-level quantification, thus enhancing the resolution of analysis for cell populations and leading to the identification of locus-specific TE markers.

MATES provides features and functionalities for single-cell TE quantification and analysis that are absent or limited in existing solutions, including the ability to handle multi-mapping TE reads. Central to MATES is a deep learning model which can analyze the distribution of background reads mapped to TE loci and their surrounding areas. This understanding of background read distribution is important for assigning reads to TE regions, thereby enhancing the clarity of genetic regulatory activities within biological systems. Distinctively, MATES achieves locus-level TE quantification for individual cells, a step beyond existing methods that typically provide subfamily level TE quantification or omit multi-mapping reads altogether. This approach leverages the inherent heterogeneity of single-cell data. By doing so, MATES allows for the exploration of locus-specific interactions between genes and TEs across different cell populations, providing insights into their contributions to broader biological processes. Moreover, MATES’ compatibility with a variety of single-cell sequencing platforms and diverse omics data types broadens its applicability and supports multi-omics analysis of the TE system. This versatility fosters the potential for understudied biological insights, enabling researchers to glean detailed information from multi-omics data. In addition, MATES brings enhanced functionality with its whole-genome TE quantification and visualization capabilities within specific cell populations. The ability to visualize whole-genome TE expressions in selected cell groups, utilizing tools like IGV, facilitates the rapid identification of potential gene-TE interactions. Complemented by an online visualization interface, MATES offers an accessible and efficient platform to analyze and interpret complex TE-related data. This interface simplifies the exploration of TE expressions and interactions, allowing researchers to intuitively navigate and interpret the genomics landscape. By integrating interactive features and user-friendly tools, MATES eases the process of identifying gene-TE interactions and their implications across different cellular contexts, thereby enriching the field of genomics and epigenetics.

The methodological characteristics of MATES have enabled several biological discoveries, which warrant further experimental validation. In the context of chemically reprogramming mouse cells to an early embryonic-like state, MATES effectively identified locus-level TE markers. Notably, markers such as MT2 and MERVL-int were enriched in the Zscan4 gene cluster, particularly flanking the *Zscan*4*c* and *Zscan*4*d* genes in 2CLC cells. This analysis also identified potential TE markers at various stages of reprogramming, illustrating the model’s capability to detect TE markers during dynamic cellular reprogramming processes. When applied to human glioblastoma data, MATES successfully identified TE locus markers specific to cancer cells and immune cells. This demonstrates its potential in pinpointing target TE loci and their interactions with nearby genes, which may contribute to advancements in cancer treatment. Furthermore, MATES was applied to adult mouse brain data to analyze chromatin accessibility in TE regions. This application revealed that the ERVK TE family is highly expressed in cells such as macrophages and astrocytes, highlighting the versatility and broad applicability of the MATES framework in diverse biological contexts. Notably, employing MATES for sc-multi-omics data has uncovered the epigenetically poised state of TEs. This underscores the advantage MATES provides for multi-modal analysis, enhancing our understanding of the complex regulatory roles TEs play across different biological states and conditions.

While MATES, a deep neural network-based method, typically requires more computational resources such as GPUs than its counterparts scTE and SoloTE for quantifying TEs, its memory usage remains relatively stable (generally below 30,000M) across single-cell datasets of varying sizes. This is attributed to its batch-based model training. Additionally, MATES’s running time is not significantly impacted by the number of cells in a dataset. For datasets of approximately 2000 to 5000 cells, MATES can complete TE quantification within 200 minutes on a computer with specific hardware configurations (Intel Xeon Platinum 8160 CPU @ 2.10GHz, NVIDIA A16), excluding time for read preprocessing and mapping (Supplementary Fig.S17 a). This sublinear increase in time and memory requirements is partly due to our downsampling strategy used in model training, as not all TE instances from every cell are necessary for training. This approach allows MATES to efficiently process large-scale single-cell datasets without being constrained by memory limitations or prohibitive running times. However, the total runtime, including read preprocessing and mapping, tends to increase linearly with data size. Overall, the total running time of MATES is generally comparable to that of scTE and SoloTE (Supplementary Fig.S17 b).

Despite the advantages of using MATES, it has limitations in quantifying certain TEs, particularly long autonomously expressed transposable elements like L1HS, which often span several thousand bases, and identical copies of TEs, such as HERVK isoforms. These cases present challenges in distinguishing between loci due to their high sequence similarity and the lack of uniquely mappable flanking regions. Although MATES could potentially struggle with the quantification of these challenging long TEs, such instances are very rare and constitute only a small portion of the genome (Supplementary Fig.S18). To accommodate the flanking regions surrounding those longer TEs, MATES does provide the flexibility to adjust relevant parameters, such as the length of the coverage vector. This adjustment could, to some extent, mitigate the potential limitations arising from insufficient coverage of long TEs. Nevertheless, these adjustments are not definitive solutions, and users should remain cautious and fully aware of the inherent limitations.

In summary, the introduction of MATES can bring into focus the intricate dynamics of TEs, converting what was once considered ‘junk reads’ into a treasure trove of information that can redefine our understanding of cellular diversity and dynamics. As single-cell technologies evolve, accurate and precise TE quantification will become increasingly crucial. MATES is poised to be an important asset in this endeavor, guiding researchers through the genomic complexities to discover new facets of biology at the single-cell level via the lens of transposon.

## Methods

### Raw reads preprocessing

Raw reads (fastq files) are mapped to the genome using STAR [73] for Smart-Seq2 scRNA-seq and STAR-solo [74] for 10x scRNA-seq, and the mapped reads are segregated into unique mapping reads that align to a single locus, and multiple mapping reads that potentially map to multiple genome locations (Fig.1 a). For the detailed command, please refer to Supplementary Note 1. To further enhance usability, we incorporate an optional realignment step into our pipeline for users who have aligned BAM files but did not keep multi-mapping reads in them. This optional step will realign those multimapping reads to ensure their inclusion in downstream analyses. The realignment process includes extracting non-unique mapping reads from the raw FASTQ files and realigning them using STAR-solo with the ‘−−outFilterMultimapNmax’ option (For details please refer to ‘realign Unmapped.sh’ in GitHub). We subsequently integrate the re-assignment with the provided BAM file to generate a new BAM file that includes all uniquely mapped reads and re-assigned multi-mapping reads.

Following alignment, samtools [75] is used to separately extract unique mapping reads and multi-mapping reads from the alignment files (i.e., bam) and store them into separate bam files. The unique mapping reads are identified by checking if MAPQ equals 255, and multi-mapping reads are extracted according to the bam file attribute NH, which indicates the number of loci the reads map to. For 10x genomics sequencing data, we have an extra step to build individual bam files for each cell according to barcodes from bam files. We then use these bam files to construct two distinct read coverage vectors (*V_u_* and *V_m_*) centered on the individual TE regions. *V_u_* represents the coverage vector of unique mapping reads, while *V_m_* denotes the coverage vector of multi-mapping reads around the TE regions. This procedure is built based on pysam and pybedtools [75–78]. In this context, “coverage” refers to the count of sequencing reads mapped to a specific genomic location. These coverage vectors serve to model the expression pattern of TEs.

To comprehensively cover TE regions, our method involves constructing a coverage vector centered at the start position of each TE region (locus). Given that over 97% of TE regions are typically under 1000 base pairs (bp) in length, as illustrated in Supplementary Fig.S19 a,b, we set a maximum TE region length (*S*) of 1000 bp which is longer than 96.8% Human TEs and 97.3% Mouse TE, and it can also be adjusted by users. Accordingly, we define the coverage vector to span 2001 bp, extending 1000 bp both upstream and downstream from the TE start position. This strategy ensures that not only is the entirety of the TE region captured in most instances but also a minimum of 1000 bp of flanking genomic sequence is included for context. For the minority of TE loci exceeding 1000 bp (approximately 3%), we truncate these regions to the initial 1000 bp. This truncation is crucial to maintain uniformity in the input vectors for the neural network, ensuring that each input vector corresponds to the same number of neurons in the network’s input layer.

In our approach, we address the challenge of overlapping regions between TEs and nearby genes. To tackle the issue of double-counting reads and the risk of information leakage at the intersections of TE and gene regions, we incorporate a specific preprocessing step. This step, executed using the bedtools [77], involves trimming the TE reference against the gene reference alignment. Such trimming effectively removes overlaps between TEs and genes, reducing the likelihood of inflated TE read counts due to these intersections.

However, it’s important to recognize that this preprocessing may not always be preferable in every application. While we have applied it in our current study to ensure that the contribution of TEs to downstream analyses is accurate and not confounded by overlapping genes, in some practical scenarios, maintaining the natural structure of these regions might be more beneficial. Trimming overlaps with gene regions could unintentionally distort TE quantification, particularly in cases where TEs intersect with neighboring genes. Therefore, in real-world applications, opting out of this preprocessing step could be advantageous. By doing so, the natural intersections of genes and TEs are preserved, allowing for a more authentic representation of their influence on TE quantification. Moreover, for those who wish to use the unaltered references, this option remains available in MATES, providing flexibility to accommodate different research needs and perspectives.

To address the need for analyzing long full-length elements, we allow users to choose the TE region length (*S*) based on their data distribution. Users can specify which TE they want to analyze in the TE reference and modify the maximum length (*S*) of the TE region. The coverage vector will now be set as twice the length of *S*, extending *S* bp both upstream and downstream from the TE start position. Additionally, users can decide whether to use the first *S* bp or the last *S* bp for the entire analysis(Supplementary Fig.S19 d). An additional experiment performed on the 10X scRNA dataset (which is 3’ sequencing data) shows that using the first *S* bp provides better performance on both ARI and NMI evaluation metrics. The results are presented in Supplementary Fig.S19 e. We also provide a downstream analysis option for those interested in specific TE loci/TE families, allowing the generation of a full-length TE count matrix.

It is important to note that the flanking region is only used to estimate the multi-mapping rate and not the reads assigned to the TEs. Our TE quantifications consider the entire TE region, which may be longer than 2001 bp. Furthermore, over 97% of TEs have lengths less than 1000 bp, making our current approach applicable to the majority of TEs (Supplementary Fig.S19 a,b). For some rare TEs that span over 2001 bp, the flanking region may not be able to cover the entire TEs, and thus the estimated multi-mapping rate may not be very accurate due to the incomplete coverage. However, those TEs are very rare (*<* 3%), and estimation based on partial coverage may not diverge significantly from the complete estimation. These options ensure that our tool can accommodate different user preferences and improve the accuracy of TE expression quantification.

### Unique-dominant and Multi-dominant Bins

The key of our method is the accurate attribution of multi-mapping reads to specific TE loci, achieved by estimating the multi-mapping reads ratio *α_i_* for each TE locus *i*. This estimation relies on the distribution of reads (i.e., read coverage vectors) around each TE locus. In a given cellular context, we choose TE regions that encompass both uniquely mapped and multi-mapped reads from the same cell as our training samples. Subsequently, we divide the TE locus (region) into small bins of a length *W* (a hyperparameter set to 5 base pairs by default). Given that read distribution across a region of any gene or TE locus tends to be smooth, we postulate that neighboring regions in close proximity within the same TE locus should have similar read distributions. This assumption is well-supported by the fact the coverage difference between neighboring bins within the same TE locus is close to zero as we observed in our single-cell data (Supplementary Fig.S19 c). Please be aware that this histogram is derived from TE loci predominantly composed of unique reads with confident mapping (TEs characterized solely by U bins), thereby avoiding significant multi-mapping uncertainty and ensuring reliable read coverage. We then employ this supported assumption to provide constraints for multi-mapping read assignments, thus incorporating it as a part of the loss term for training our deep neural network.

To ascertain the proportions of multi-dominant (M) and unique-dominant (U) bins within each TE region, we apply a *W* bp slicing window. M and U bins are determined by the percentage of unique and multi-mapping reads within each bin. A bin is categorized as an M-bin if the percentage of multi-mapping reads exceeds a threshold *P* ; otherwise, it is deemed a U-bin. Selection of the hyperparameters, the bin size (*W*) and proportion threshold (*P*), is dependent on the data’s multi-mapping rates. For data with high multi-mapping rates, we opt for a smaller bin size and a higher threshold to enhance accuracy. Conversely, for low multi-mapping rate data, we prefer a larger bin size and a lower threshold to acquire more training samples (please see Supplementary Fig.S19 d,f for the selection of those parameters). The motivation to segregate bins into U and M categories stems from the desire to utilize the more reliable U bins (where unique reads are assigned confidently) to constrain the estimation of the multi-mapping rate within a specific TE locus (Fig.1 b).

### Estimating the multi-mapping rate for each TE locus via a deep neural network

TE loci are divided into three specific groups based on the presence of unique-dominant (U) and multi-dominant (M) bins. The groups comprise TE regions that contain both M and U bins; TE regions composed solely of M bins; and TE regions that consist only of U bins. Our model utilizes TE regions from the first group—those containing both M and U bins—as training samples. The presence of both types of bins in these regions is necessary for calculating the loss used in optimizing our neural network. It’s important to highlight, however, that certain TE families may have a higher degree of repetitiveness, leading to a higher occurrence of M bins in their instances. Consequently, our training dataset, curated from single-cell data, might lean towards an overrepresentation of these specific TE families. To counteract this potential imbalance, we have implemented a downsampling strategy, where we limited training samples for each TE family not exceeding 2500 inputs (Larger values are recommended to enhance TE quantification performance, although this will result in extended training time). This approach aims to decrease the overrepresentation of certain TE families, thereby fostering a more balanced representation within our model. We also removed TE families with too little training sample (less than 50 samples) to focus on more representative TE families. Such balanced representation is instrumental in enhancing the model’s overall performance and accuracy. Moreover, our downsampling strategy results in the size of the training sample being more dependent on the number of expressed TE families in the dataset than on the total number of cells. This method helps in maintaining both training time and memory usage at a steady level, regardless of the cell count.

Next, we employ coverage vectors constructed from unique mapping reads *V_u_* to train the AutoEncoder(AE) model [79]. Recognizing that different TE families may exhibit distinct expression patterns, we introduce TE family information into the input and embedding layers to account for such heterogeneity. This is accomplished by encoding the TE family information of each TE region using one-hot encoding. The AE model minimizes the Mean Squared Error loss function (*L*_1_) to minimize the discrepancy between the input vector and the reconstructed vector, resulting in a more meaningful reduced embedding of the TE locus (Fig.1 c).

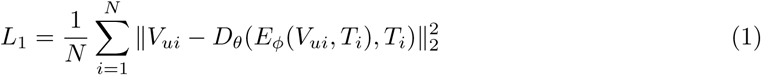

Here N represents the number of training samples, *V_ui_* represents the unique mapping reads coverage vector of training sample TE locus *i*, while the *T_i_* is its one-hot encoded TE family information for the specific TE locus. *E_θ_* and *D_ϕ_* represents encoding function and decoding function respectively. Subsequently, the trained AE model generates embeddings to represent the read distribution of the TE loci in the latent space. These embeddings are combined with the TE family information of the corresponding TE loci and utilized as the input for a Multilayer perceptron (MLP) regressor [80] (Fig.1 c). For the MLP, we define the loss function (*L*_2_) to minimize the read coverage difference between each M-bin (*j*) and its nearest neighboring U-bins (*N_j_*). The loss function (*L*_2_) is defined as follows:

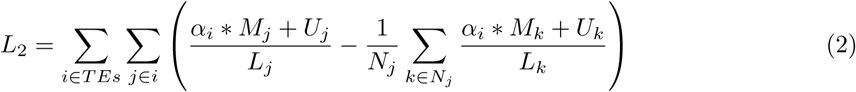

Here, *M_j_* and *U_j_* represent the total number of multi-mapping reads and unique mapping reads falling within the j-th M-bin of TE locus *i*, while *L_j_* represents the number of covered base pairs in this region. Similarly, *M_k_* and *U_k_* represent the total depth of the *k*-th U bin among the *N_j_* nearest neighboring U bins of the *j*-th M bin, and *L_k_* represents the number of covered base pairs in that *k* − *th* bin. The output *α_i_* represents the probability that a multi-mapping read falls into this TE locus. The grand loss to train the entire model is the sum of both loss terms *L_grand_* = *L*_1_ + *λ* ∗ *L*_2_, where *λ* is a hyperparameter.

### TE quantification and characterization at the single-cell level

In our study, we employ a neural network framework that integrates an Autoencoder (AE) with a Multi-layer Perceptron (MLP) regressor. This hybrid model can assign multi-mapping reads to specific TE loci, a crucial step in quantifying TE expression in individual cells. The process begins by feeding the model a read coverage vector. This vector, which is derived from unique-mapping reads, is specific to a given TE locus *i*, denoted as *V_ui_*, accompanying this vector is the TE family information associated with the TE instance. Our model, leveraging its neural network capabilities, computes a multi-mapping probability, *α_i_*. This probability reflects the likelihood of multi-mapping reads being associated with the TE locus *i*. Subsequently, this computed probability is utilized to estimate the count of multi-mapping reads for the TE locus in question. It achieves this by multiplying *α_i_*, with the aggregate of all potential multi-mapping reads within that particular TE locus *i*. The outcome of this multiplication is an estimated count of multi-mapping reads assigned to the specific TE locus. The total read number of a given cell at TE locus i is then calculated by *UR_i_* + ⌈*α_i_* × *MR_i_*⌉, where *UR_i_* is the unique read number and the *MR_i_* is the multi-read number at the TE locus i respectively.

To facilitate a comprehensive understanding of TE expression at the cellular level, we construct a cell-by-TE expression matrix. This matrix is a confluence of two critical components: the estimated counts of multi-mapping read (as determined by our neural network model) and the counts of uniquely mapped read corresponding to the same TE locus. We have meticulously designed this matrix to integrate seamlessly with the conventional cell-by-gene expression matrices. This compatibility is pivotal for enabling effective downstream analyses. One such downstream analysis is cell clustering, which is significantly enhanced by incorporating our cell-by-TE expression matrix. By using this matrix for cell clustering, we can underscore the effectiveness and precision of our TE quantification method. An illustrative example of this process and its outcomes is presented in Fig.1 d.

After identifying distinct cell populations in the single-cell data with TE quantification from MATES, we further employ the biomarker calling method from SCANPY [81], which systematically identifies signature genes and TEs that show higher expression in specific cell populations compared to other cells. Its primary function is to distinguish signature genes and TEs unique to each cell population.

MATES enhances its functionality by producing TE reads alignment files in BigWig format. This process involves a series of steps. Initially, coverage vectors are recalculated for each TE locus, denoted as *i*. This recalculation is done separately for unique reads and multi-mapping reads, using a comprehensive TE reference that includes all TEs, without any trimming or exclusion of TE loci overlapping with genes. Subsequently, at each base pair within a TE locus, the coverage from multi-mapping reads is adjusted. This adjustment involves multiplying the read coverage of the multi-mapping reads at each specific base pair by the locus-specific multi-mapping ratio, *α_i_*. The final read coverage at each base pair is then determined by summing the coverage from unique reads and the adjusted coverage from multi-mapping reads. The generated bigwig files include all reads assigned probabilistically to TE regions by our neural network and are compatible with genomic visualization tools such as IGV [82] (see Fig.1 e), facilitating genome-wide TE examinations and visualizations. To provide a comprehensive exploration of the outputs from MATES, including cell clusters, gene biomarkers, TE biomarkers, and the probabilistic assignment of reads across the genome, we have developed an interactive MATES web server, available at https://mates.cellcycle.org. This platform allows users to interact with and analyze the data generated by our method, offering a resource for detailed investigation of TEs and genes across different cell populations.

### Different modes for TE quantification

MATES operates in three modes: ‘exclusive’, ‘inclusive’, and ‘intronic’. In the ‘exclusive’ mode, any TE region overlapping with gene regions is excluded from quantification to avoid information leakage from the overlapping genes that could introduce bias towards the cell clustering evaluation. The ‘inclusive’ mode utilizes all TE regions, including those overlapping with genes. Under this mode, all reads from the intronic TEs are used for the TE quantification. Intronic TEs constitute a significant portion of all TEs, and excluding them might lead to loss of crucial information.

The ‘intronic’ mode addresses the critical issue of distinguishing whether reads assigned to intronic TE loci originate from TE expression or from the TE-residing genes. For scRNA-seq data, if the reads assigned to the intronic TEs originated from the TE-residing gene, those reads are most likely from unspliced RNAs of the gene that contains the introns. Therefore, identifying all unspliced reads assigned to the intronic TEs can, to some extent, correct the TE quantification. In the ‘intronic’ mode, the method for intronic TE quantification involves the following steps. First, the total number of reads (*N*) aligned to intronic TEs is quantified. Then, using Velocyto [83], the number of unspliced reads (*s*) mapped to each intronic region is identified with the command ‘velocyto run -b path to bam annotation.gtf –dump p1’. Finally, the final intronic TE counts (*N ^′^*) are calculated as *N ^′^* = *N* − *s*.

Since Velocyto does not provide detailed information on which specific reads are unspliced, custom code adapted from the Velocyto (version 0.17) source was added to parse the results and categorize the reads. This allows for accurate alignment of unspliced reads to intronic TE regions and adjustment of the final quantification accordingly. Detailed implementation can be found on our GitHub page (https://github.com/mcgilldinglab/MATES).

The model with intronic TE correction was applied to the 2CLC single-cell RNA-seq dataset. Supplementary Fig.S20 a shows that the contribution of reads from nearby genes, detected via unspliced RNAs, is marginal. This marginal contribution explains why the cell representation and clustering results using all reads are only slightly better than those using the corrected reads. The cell representation and clustering using the corrected intronic TE reads are similar to those using the raw intronic TE reads (which include potential genes from overlapping regions) (Supplementary Fig.S20 b), with ARI values of 0.364 vs. 0.390 and NMI values of 0.490 vs. 0.524.

These different modes of MATES ensure that the method can be adapted to different scenarios, including accounting for the complexities associated with intronic TEs, thus providing more accurate and robust quantification.

### MATES application to single-cell data across different modalities

In order to demonstrate the broad applicability of MATES, we evaluated this approach on single-cell datasets across diverse modalities and multiple platforms. These include 10x scRNA-seq, Smart-Seq2 scRNA-seq, 10x scATAC-seq, and 10x multiome. We implemented a consistent processing sequence for these datasets, regardless of their platform or modality.

For single-cell RNA-seq datasets derived from platforms like 10x and Smart-Seq2, we adhered to the aforementioned procedure. When conducting single-cell ATAC-seq analysis, we chose STAR-solo for read mapping due to its ability to handle multi-mapping reads, unlike traditional single-cell ATAC-seq read mapping tools such as CellRanger ATAC which discard these multi-mapping reads [84]. We then applied MATES to the bam files generated by STAR-solo, leading to the creation of a cell-by- TE chromatin accessibility matrix. Concurrently, we generated the conventional cell-by-peak count matrix using our chosen tool, in this case, the CellRanger ATAC toolset. Nonetheless, users have the flexibility to perform conventional peak quantification using any tool of their preference. This matrix was integrated with the cell-by-TE matrix derived from MATES via concatenation, enabling subsequent downstream analyses like cell clustering. In cases where users engage with other single-cell modalities not discussed in this study, such as single-cell DNA methylation profiling, they can utilize MATES to generate the cell-by-TE matrix for downstream analysis. However, the extraction of cell-by-gene or cell-by-peak matrices connected to the cells can be achieved through established pipelines, which fall outside the purview of our TE-focused study.

For the processing of multi-omics datasets, we employed a multi-step strategy. Initially, we performed independent alignments for each modality, using the STAR-Solo alignment tool and adhering to the specific considerations previously mentioned. Using the MATES method, we subsequently quantified individual TE for each modality. Following this, we consolidated the TE expressions from all modalities with their respective gene/peak expressions, quantified through conventional methods such as Cell Ranger. We then concatenated the cell-by-TE matrices across modalities or integrated them using other multi-modal data integration methods, like Seurat [85]. The resulting integrated multi-omics TE matrices offered by MATES provide a comprehensive representation of the cells, leading to enhanced cell clustering and identification accuracy. Furthermore, multi-omics gene and TE quantification allow us to explore the roles of TEs in the identified cell populations from different modality perspectives, thus yielding deeper biological insights.

### Evaluate the similarity between TE-based clustering and gene-based clustering results

To evaluate the biological conservation of TE quantification, we performed clustering using TE expressions and calculated ARI (or NMI) scores against clustering results based on gene expressions. To assess the similarity between TE clusters and gene clusters, we used block permutation [86] to build the background distribution and calculate the p-value, statistically examining their similarity. In conducting block permutation, the predicted labels are divided into blocks with a certain number of cells (block size), and the order of the blocks is permuted to form the randomized prediction results. This permutation is conducted multiple times to form the random distribution to calculate p-value by examining whether ARI between TE-clusters and gene-clusters is achieved by chance or not. Empirically, we performed the block permutation 1,000,000 times on the TE cluster labels with a block size of 300 to build the background distribution. This random background distribution was then used to calculate permutation-based p-values, determining the statistical significance of similarity (based on ARIs and NMIs) between TE-based and gene-based clustering results for each method.

### MATES comparison with existing TE quantification methods on bulk data

We compared the predictions from MATES against those from established bulk data TE quantification tools, TEtranscripts and Telescope, to showcase the effectiveness of our method. Both TEtranscripts and Telescope are EM algorithm-based tools capable of analyzing a wide range of transposable elements. To compare MATES with these methods, we employed pseudo-bulk datasets to assess the applicability and performance of our tool.

MATES was designed specifically for single-cell data, leveraging the cellular heterogeneity. Training MATES requires a large volume of cell profiles, which the scale of bulk data usually cannot provide. To facilitate meaningful comparisons at the bulk level, we trained MATES using single-cell datasets and aggregated single-cell TE expressions by adding together the expression levels in each cell to form a surrogate pseudo-bulk-level TE expressions for each sample. To run benchmarking methods, we generated pseudo-bulk datasets from single-cell data obtained from the 10X platform, which produces one BAM file per sample, similar to bulk data. The key difference is that 10X BAM files include CB/CR fields, representing cell barcodes, indicating which cell each read originates from. By ignoring the CB/CR fields, we treated the single-cell BAM files as bulk BAM files and directly applied bulk TE quantification tools on them.

To further evaluate MATES’s accuracy and efficiency, we calculated the coefficient of determination (*R*^2^) between pseudo-bulk-level TE expression from MATES and the bulk TE quantification tool and used F-test to evaluate the significance of the score. We specifically included well-defined TEs for the RNA analysis and all repetitive elements for the ATAC analysis to ensure comprehensive coverage. We then created a scatter plot where each dot represents a TE at the subfamily level. The y-axis of the plot corresponds to the TE expression levels quantified by the bulk TE quantification tool, while the x-axis represents the pseudo-bulk expression levels quantified by MATES. To quantify the correlation between these two methods, we fitted a linear regression line to the scatter plot data points. The *R*^2^ value derived from this regression analysis indicates the degree of correlation, thereby assessing MATES’s performance in approximating bulk-level TE expression from single-cell data. This detailed comparison allowed us to rigorously evaluate the effectiveness of MATES against established bulk quantification tools.

### Method benchmarking design

To benchmark the accuracy of our TE quantification method, we focused on evaluating its impact on cell clustering performance within single-cell datasets. The rationale behind this approach is based on the hypothesis that more accurate TE quantification leads to enhanced cell clustering and identification. This enhancement in cell clustering can be quantitatively assessed using two key metrics: ARI and NMI. Both metrics are crucial in single-cell research, as they provide a means to compare clustering results against known cell type labels, offering a clear indication of clustering accuracy. Our benchmarking process involved comparing the performance of our method with other state-of-the-art TE quantification methods, scTE and SoloTE. SoloTE is known for its ability to quantify unique read TE expression at the locus level and multi-mapping read TE expression at the subfamily level, while scTE quantifies both at the subfamily level. To ensure a consistent and fair comparison, we normalized the data by aggregating locus-level TE expressions to subfamily levels across all methods. This standardization is vital for making meaningful comparisons between the methodologies. The effectiveness of each method in improving cell clustering performance was statistically evaluated using the one-sided Student’s t-test. This test determines whether the improvements in ARI and NMI scores with a particular TE quantification method are statistically significant compared to others.

In our study, we sought to showcase the added value of TEs as an informative component in addition to traditional gene expression analysis. To achieve this, we conducted cell clustering analyses under different scenarios: using only gene expression, combining gene and TE expressions, and using TE expression alone. The clustering based on gene expression alone established a baseline for comparing the enhanced performance achieved when TE data was incorporated using the three methods. Additionally, to specifically understand the impact of TE expression on clustering performance, we performed clustering analyses solely based on TE expression. This allowed us to isolate the effect of TE data from the gene expression influence. To create a foundational comparison point, we utilized a unique-mapping reads-only approach for TE quantification. This approach, which omits multi-mapping reads, provided a clearer perspective on the efficacy of each method’s TE quantification process. Furthermore, we evaluated the effectiveness of TE expression quantification when relying exclusively on multi-mapping reads. This was crucial to demonstrate the capability of our method, particularly in handling the complexities of multi-mapping scenarios through context-aware expression analysis. MATES underwent extensive testing across various single-cell modalities and platforms, highlighting its versatility and broad applicability in different research contexts.

### Validation with nanopore and pacbio long-read sequencing data

#### Validation using the scNanoRNAseq data

To validate the accuracy of our TE quantification model, we employed a melanoma brain metastasis nanopore long-read sequencing (scNanoRNAseq) data [66]. The long-read scNanoRNAseq data can lead to lower multimapping rate, thus serving as a proxy ground truth for TE expression. This dataset contains long-read data with an average length of 937 base pairs (bp). This is around 4x longer than the corresponding short-read data of 222 bp obtained from 10x scRNA-seq, which came from the same set of cells in the study. The longer read length of nanopore sequencing is advantageous for TE quantification, as it allows for more accurate read alignment [68] (i.e., reduced multi-mapping rate ∼ 1% in nanopore sequencing compared to over 12% in 10x shortread sequencing from the same study), thereby enhancing the reliability of TE quantification. This level of precision from long-read data serves as the ground truth for our validation of short-read TE quantification by MATES. We applied both our method and other benchmarked methods to the 10x short-read sequencing data from the same study, corresponding to the same cell sets analyzed via the long-read sequencing approach. The focus was on comparing the TE expression levels quantified by MATES, scTE, and SoloTE from the short-read data against the long-read data, considered as the ‘ground truth’. For this comparison, we used the *R*^2^ as the metric to assess the accuracy of TE quantification by each method. A higher *R*^2^ value, indicating greater agreement with the long-read ground truth, would suggest a more precise TE quantification from the matched short-read data by each method. To enhance the accuracy of our correlation calculations and accurately reflect the relationship between short-read and long-read expressions, we included only well defined TE families (DNA, LTR, RC, Retroposon, SINE, and LINE). This was to ensure a focused and accurate analysis of transposable elements and the *R*^2^ values truly represented the biological data, free from distortions caused by non-expressive or anomalous readings. This approach allowed us to evaluate the accuracy of TE quantification methods using the nanopore long-read sequencing data as a benchmark, providing a validation of our model against other existing techniques.

### Validation using the Pacbio data

To further strengthen our validation, we also utilized PacBio long-read data from the postnatal mouse brain. The PacBio long-read data was aligned using STAR following the command provided in [69], and cell barcodes were deconvoluted using Scisorseqr [69]. The processed data contains 7,896 cell barcodes with reads of an average length of 1,164 bp and a variance of 521.7 bp. Similarly, the dataset also contains corresponding 10X short-read data of 91 bp, which came from the same set of cells in the study. We mapped these reads to the transcriptome, utilizing transcriptome annotations to guide the mapping process. We employed a strategy similar to the benchmarking experiments with the scNanoRNAseq data, comparing MATES against scTE and SoloTE by running on the short-read data. We used TE quantification from the PacBio long-read data as the ground truth and *R*^2^ as the metric to assess the accuracy of TE quantification by each method. Notably, we included all TEs, including repetitive ones, to comprehensively evaluate the performance of each method using the PacBio dataset.

### Validation with controlled simulation

To address the limitations of nanopore sequencing in capturing short TEs, we randomly selected 6 subfamilies from the Alu family in the SINE class and 15 subfamilies from the Simple Repeats family to conduct controlled simulations. For each TE region belonging to these subfamilies within the human genome, we simulated reads, extending the simulation to include 2000 base pairs surrounding each TE region—1000 base pairs upstream and 1000 base pairs downstream. Using Polyester [87], a tool that simulates reads directly from specified genomic regions and produces reads, we randomly simulated between 1 and 1000 reads for each locus. In our simulation of Alu family, six TEs from the Alu family (AluYa8, AluYc, AluYc3, AluYe6, AluYf1, and AluYh3a3) were randomly selected, and a total of 1,533,859 reads were simulated. The simulated reads were aligned using STARsolo. After alignment, 90.36% of reads were uniquely mapped, 7.19% were mapped to multiple loci, and 1.64% were unmapped. The simply repeats simulation follows the same strategy as Alu family simulation. The simulated reads were aligned using STARsolo. After alignment, 96.60% of reads were uniquely mapped, 3.17% were mapped to multiple loci, and 0.23% were unmapped. We assigned a pseudo cell barcode to the simulated reads to produce Smart-seq2 formatted data [87]. Then we merged them with all other reads from actual single-cell RNA-seq data. The MATES model, trained on the original scRNA-seq data, was then used to process and quantify the TE levels in the combined dataset (simulated reads plus all reads from the original data). To ensure an accurate ground truth reads from the original data that were also mapped to the selected TE subfamilies were removed. The quantifications by MATES were compared against the simulated ground truth, resulting in an *R*^2^ value. Since SoloTE requires cell barcodes and unique molecular identifiers (UMIs) to identify cells (and does not support Smart-seq2 formatted data), we focused on comparing the subfamily expression levels quantified by scTE and MATES.

Repeats can form different TE isoforms through alternative splicing. To investigate the performance of MATES in handling repeats from different isoforms, we conducted a series of targeted simulation studies using human genome. We selected two HERVK-int regions, TE1 and TE2, to represent HERVK with and without the env deletion. TE1 (*chr*3|75551382|75558968| + |*HERV K* − *int*|*ERV K*, 7586 bp) acts as HERVK without the deletion, while TE2 (*chr*4|3978357|3985909| − |*HERV K* −*int*|*ERV K*, 7542 bp) represents HERVK with a 292 bp deletion (Supplementary Fig.S15 c). We simulated 5000 reads for TE1 and 2000 reads for TE2 (excluding the 292 bp deletion region), including 1000 bp flanking both sides. The simulation was performed using Polyester[87], with STAR-solo used for alignment. For the TE1 locus, out of the 5000 simulated reads generated, 4126 reads were successfully mapped to the TE1 locus using STARsolo. The remaining 874 reads were located in TE1’s flanking 1000 bp regions on both side. For the TE2 locus, 1282 out of 2000 simulated reads were mapped to the TE2 region. In our analysis, we used only the reads located in the TE regions as the ground-truth numbers (TE1: 4126, and TE2: 1282) to evaluate the effectiveness of MATES. We then applied the MATES model, pre-trained on the human scRNA-seq data from a multi-omics dataset described earlier, to predict the number of counts in the TE1 and TE2 regions.

To directly evaluate the accuracy of locus-level TE quantification, we conducted additional bench-marking by simulating full-length short reads from scNanoRNA-seq long-read data [66]. Specifically, we generated 100 bp short reads from the scNanoRNA-seq long-read data using SeqKit (v2.8.2) [88] with a step size of 100 bp. This approach aimed to replicate scenarios involving shorter reads, which typically increase the likelihood of multi-mapping issues while preserving the biological context provided by the long-read data. This strategy enabled a more precise assessment of the effectiveness of utilizing nearby region information to assist with multi-mapping read assignments. The locus-level TE quantification from different methods was then evaluated against the proxy ground truth provided by the nanopore long-read sequencing data. As previously mentioned, scTE does not yet support locus-level TE quantification, and SoloTE does not natively support full-length Smart-seq-like formatted data (such as those simulated from long-read data). Therefore, we benchmarked the locus-level TE quantification performance of MATES and SoloTE (using the same strategy as in the SoloTE tool) on the simulated data.

## Data availability

The TE reference data was sourced from the Repeat Masker website. Specifically, the Mouse TE reference is available at https://www.repeatmasker.org/species/mm.html, and the Human TE reference can be accessed at https://www.repeatmasker.org/species/hg.html. In this study, we utilized a range of publicly accessible benchmark datasets that cover scRNA-seq, scATAC-seq, and multi-omics datasets across various biological contexts. The scRNA-seq 10x mouse chemical reprogramming dataset (GSE114952, https://www.ncbi.nlm.nih.gov/geo/query/acc.cgi?acc=GSE11495) is detailed in Zhao et al 2018 [25], and the Smart-Seq2 human glioblastoma dataset (GSE84465, https://www.ncbi.nlm.nih.gov/geo/query/acc.cgi?acc=GSE84465) is described in Darmanis et al. 2017 [33]. Both datasets provided cell type annotations. The scATAC-seq 10x 5k adult mouse brain cell dataset [47] is available at https://support.10xgenomics.com/single-cell-atac/datasets/1.2.0/atac_v1_adult_brain_fresh_5k, and we generated the cell type labels following the workflow provided in the Signac tutorial (https://stuartlab.org/signac/articles/pbmc_vignette). The 10x Multi-Omics 10k PBMC dataset [61] is accessible at https://support.10xgenomics.com/single-cell-multiome-atac-gex/datasets/1.0.0/pbmc_granulocyte_sorted_10k, and we generated cell type labels following the workflow described by Hao et al. [89]). Additionally, the study utilized mRNAs sequenced by single-cell nanopore (GSE212945, https://www.ncbi.nlm.nih.gov/geo/query/acc.cgi?acc=GSE212945) as detailed in Shiau et al. 2023 [66] and postnatal mouse brain Pacbio long read data [69] (GSE158450, https://www.ncbi.nlm.nih.gov/geo/query/acc.cgi?acc=GSM4800803 for 10X and https://www.ncbi.nlm.nih.gov/geo/query/acc.cgi?acc=GSM4800807 for Pacbio). The Drosophila melanogaster dataset is published by Zhu et al 2022 [45] (GSE168553, https://www.ncbi.nlm.nih.gov/geo/query/acc.cgi?acc=GSM5145863) The Arabidopsis thaliana dataset is published by Zhai et al 2021 [44](GSE156991, https://www.ncbi.nlm.nih.gov/geo/query/acc.cgi?acc=GSE156991). Source data are provided with this paper.

## Software availability

MATES, developed as a Python-based tool, is publicly accessible and comes with comprehensive documentation. It can be found at https://github.com/mcgilldinglab/MATES. Additionally, for enhanced user interaction, especially in visualizing TE quantifications and cell clustering, a user-friendly web interface has been established. This can be accessed at https://mates.cellcycle.org.

## Acknowledgments

This work is supported by grants from the Canadian Institutes of Health Research (CIHR) [PJT-180505 to J.D]; the Funds de recherche du Québec - Santé (FRQS) [295298 to J.D., 295299 to J.D.]; the Natural Sciences and Engineering Research Council of Canada (NSERC) [RGPIN2022-04399 to J.D.]; and the Meakins-Christie Chair in Respiratory Research [to J.D.]; and Tao P. Wu is a CPRIT scholar for cancer research, supported by CPRIT award (RR180072). We thank MyCellome LLC for supporting the visualization of MATES.

## Author contributions

J.D. and T.P.W. conceived and designed the study, developed the methodology, and planned the experiments. R.W. and Y.Z. contributed to design, implementation of the methodology, and conducted the experiments. J.D., T.P.W., R.W., Y.Z. and Z.Z. participated in data collection and the analysis of computation experiments. X.Z. contributed to the analysis of computation experiments. All authors (J.D., T.P.W., R.W., Y.Z., Z.Z., K.S., E.W., X.Z.) contributed to writing and revising the manuscript. Each author has read and approved the final manuscript for publication.

