## Supplementary Figures for "MATES: A Deep Learning-Based Model for Locus-specific Quantification of Transposable Elements in Single Cell"

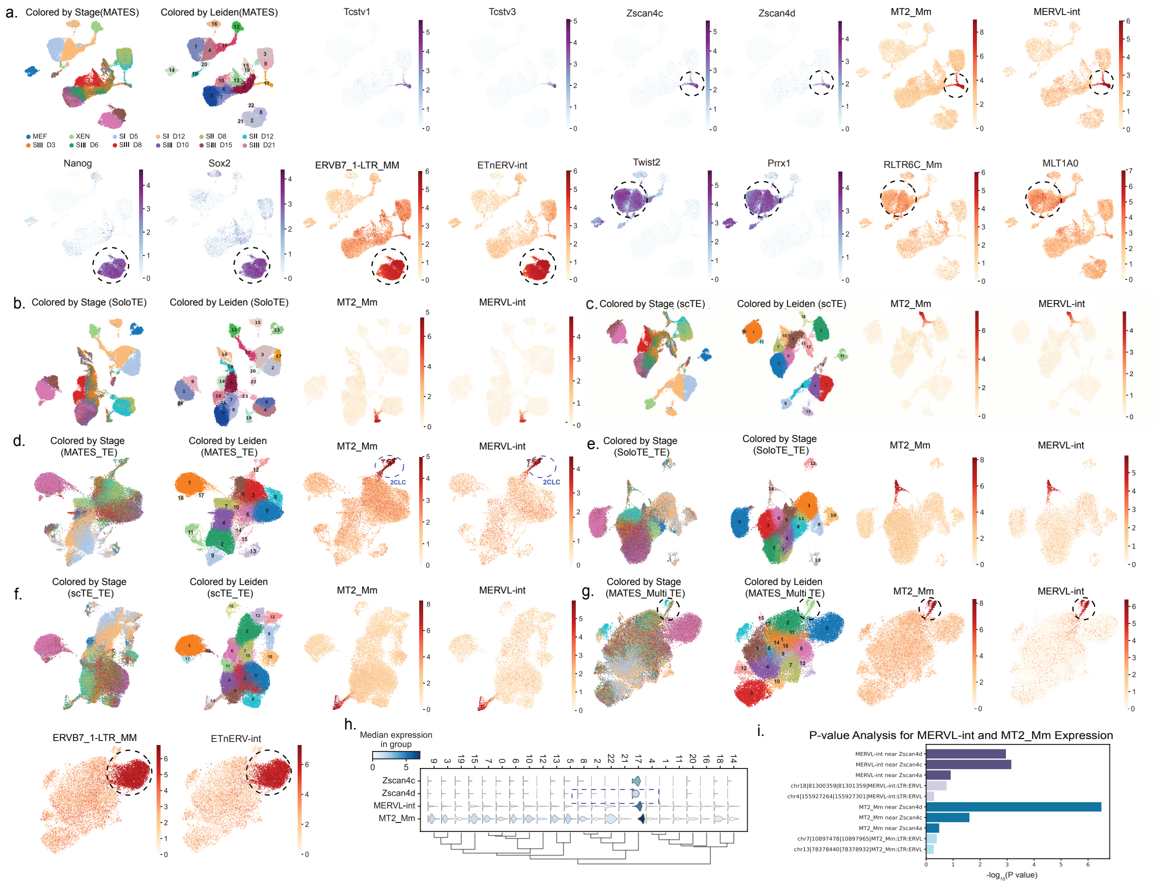

**Fig. S1 Additional analysis of chemical reprogramming dataset with 10X single-cell RNA platform.** Using MATES, multi-mapping reads from the single-cell RNA dataset were retrieved and assigned to the TE instance. Single-cell embedding was based on gene and/or TE expression. (a) UMAP visualization of cell clustering and gene or TE markers. The type of embedding is indicated in each UMAP title's parentheses. "MATES" denotes gene and TE expression determined by MATES. The MATES UMAP is colored by stages and by Leiden clustering. 2C-like gene markers (Tcstv1, Tcstv3, Zscan4c, Zscan4d) and TE markers (MT2_Mm, MERVL-int) are highlighted on the MATES UMAP. Stem cell gene markers (Nanog, SOX2) and TE markers (ERVB7_1-LTR_MM, ETnERV-int) are also shown. Additionally, MATES identified stage-specific TE markers, RLTR6C_Mm and MLT1A0, which are enriched in stage I, as indicated by Twist2 and Prrx1. (b) UMAP visualization of cell clustering and TE markers quantified by SoloTE. The type of embedding is indicated in each UMAP title's parentheses. (c) UMAP visualization of cell clustering and TE markers quantified by scTE. The type of embedding is indicated in each UMAP title's parentheses. (d)-(f) UMAP visualization based solely on TE quantification by (d)MATES, (e)SoloTE and (f)scTE, clustering cells by transposon element expression and highlighting TE markers of 2CLCs (MT2_Mm and MERVL-int). (g) UMAP visualization of multiple mapping read quantification, with stages and Leiden clusters highlighted. Key markers include MT2_Mm, MERVL-int, ERVB7_1-LTR_MM and ETnERV-int. (h) Violin plot shows the TE expression levels of 2CLC stage cells across different clusters. (i) Bar plots display read enrichment for MT2_Mm and MERVL-int at *Zscan4c*/*Zscan4d* loci compared to random MT2_Mm and MERVL-int loci, with bars indicating $-log_{10}$ p-values, derived from locus expression fitted to a negative binomial distribution, to highlight the statistical significance at these loci.

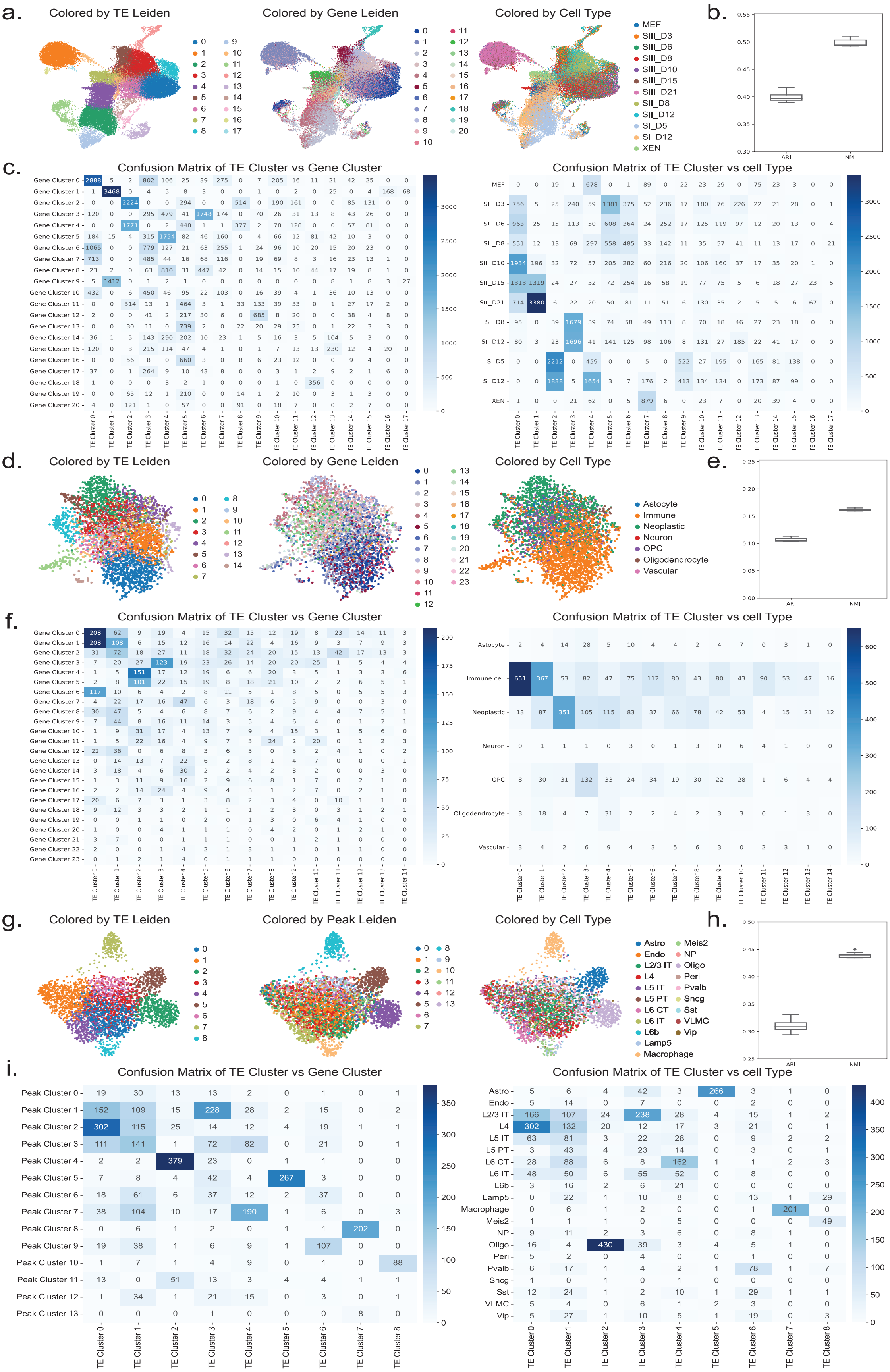


**Fig. S2 Clustering Comparison between TE and Gene quantification across different datasets.**

**Fig. S2** (a)-(c) the analysis results for 10X scRNA-seq data. (a). UMAP visualizations of cell clustering based on only the TE expression, colored by Leiden clusters, gene expression clusters, and cell type. (b). Boxplot analyses of ARI ($P < 1 \times 10^{-6}$) and NMI ($P < 1 \times 10^{-6}$) scores, comparing TE clusters to gene clusters. (c). Quantitative analyses including confusion matrices that compare TE clusters to gene expression clusters, illustrating the overlap and correspondence between clusters identified by TE expression (Leiden clusters) and gene expression clusters. Additionally, confusion matrices compare TE clusters to cell types, highlighting the overlap and correspondence between clusters identified by TE expression (Leiden clusters) and cell types. (d)-(f) the analysis results for Smart-seq scRNA-seq data. (d). UMAP visualizations of cell clustering based on TE expression, colored by Leiden clusters, gene expression clusters, and cell type. (e). Boxplot analyses of ARI ($P = 1.03 \times 10^{-2}$) and NMI ($P = 7.60 \times 10^{-4}$) scores, comparing TE clusters to gene clusters. (f). Quantitative analyses including confusion matrices that compare TE clusters to gene expression clusters, illustrating the overlap and correspondence between clusters identified by TE expression (Leiden clusters) and gene expression clusters. Additionally, confusion matrices compare TE clusters to cell types, highlighting the overlap and correspondence between clusters identified by TE expression (Leiden clusters) and cell types. (g)-(i) the analysis results for 10X scATAC-seq data. (g). UMAP visualizations of cell clustering based on TE expression, colored by Leiden clusters, peak expression clusters, and cell type. (h). Boxplot analyses of ARI ($P = 5.60 \times 10^{-5}$) and NMI ($P = 4.60 \times 10^{-6}$) scores, comparing TE clusters to gene clusters. (i). Quantitative analyses including confusion matrices that compare TE clusters to peak expression clusters, illustrating the overlap and correspondence between clusters identified by TE expression (Leiden clusters) and peak expression clusters. Additionally, confusion matrices compare TE clusters to cell types, highlighting the overlap and correspondence between clusters identified by TE expression (Leiden clusters) and cell types. The p-values of ARI and NMI are calculated against the random ARI score background built by using the block permutation strategy.

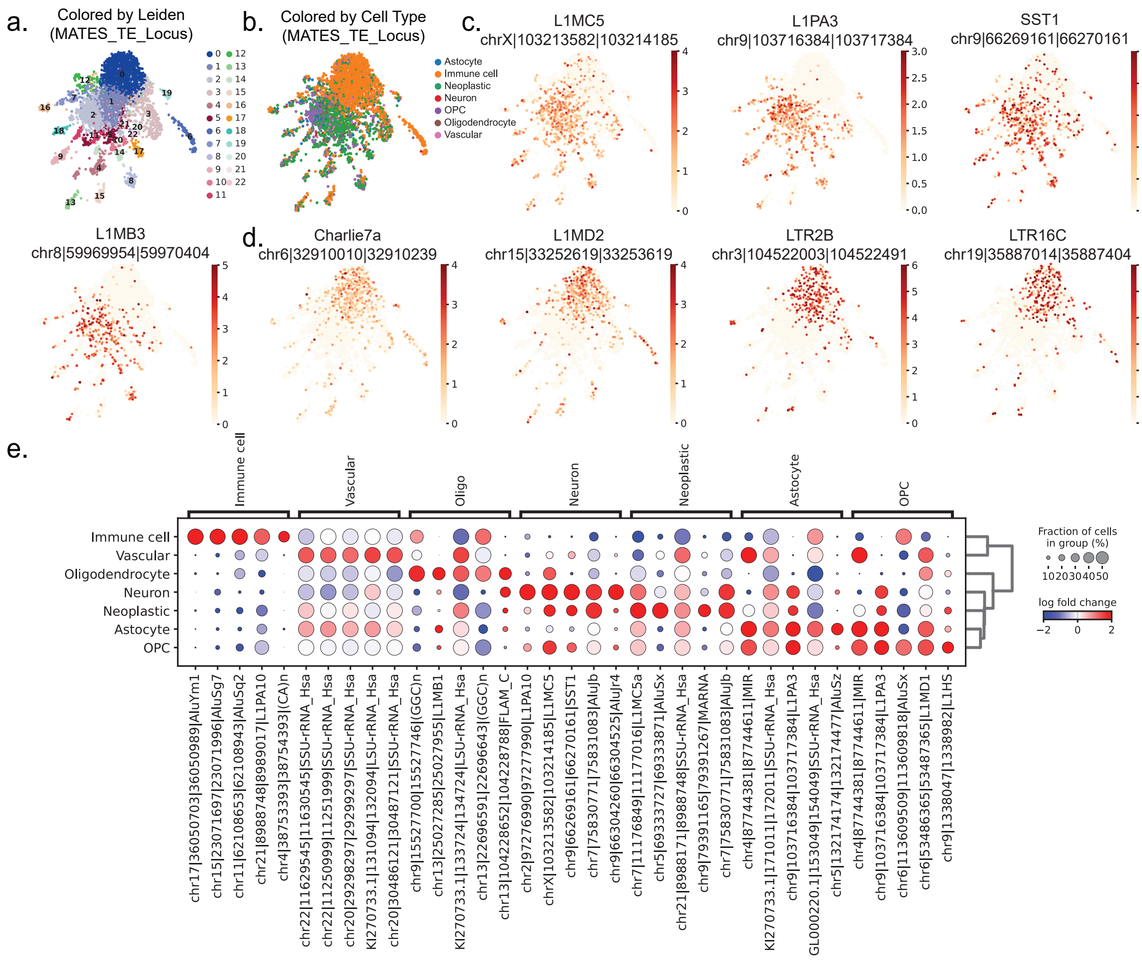


**Fig. S3 Extended Analysis of the Smart-seq2 Full-Length scRNA Dataset from Human Glioblastoma.** (a-d) UMAP visualizations illustrate cell clustering based on locus-level TE expressions in human glioblastoma cells. Panel (a) displays clusters colored according to the Leiden algorithm, while panel (b) differentiates them by cell type. Panel (c) features locus-level TE markers specific to Immune cells (including L1MC5, L1PA3, SST1, and L1MB3). Panel (d) shows locus-level TE markers for Neoplastic cells (such as Charlie7a, L1MD2, LTR2B, and LTR16C), highlighted on the UMAP. (e)A dot plot demonstrates the marker TE locus for each cell type, as identified by the MATES methodology.

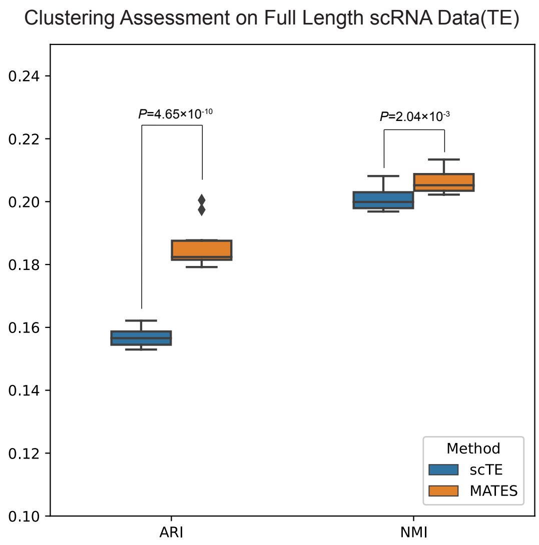


**Fig. S4 Clustering Performance Comparison for Full Length scRNA-seq.** TE Expression Quantified by Different Methods. This figure illustrates the ARI and NMI scores for TE clustering performance on Smart-seq2 by scTE and MATES respectively. The p-value was calculated using the one-sided Student's t-test.

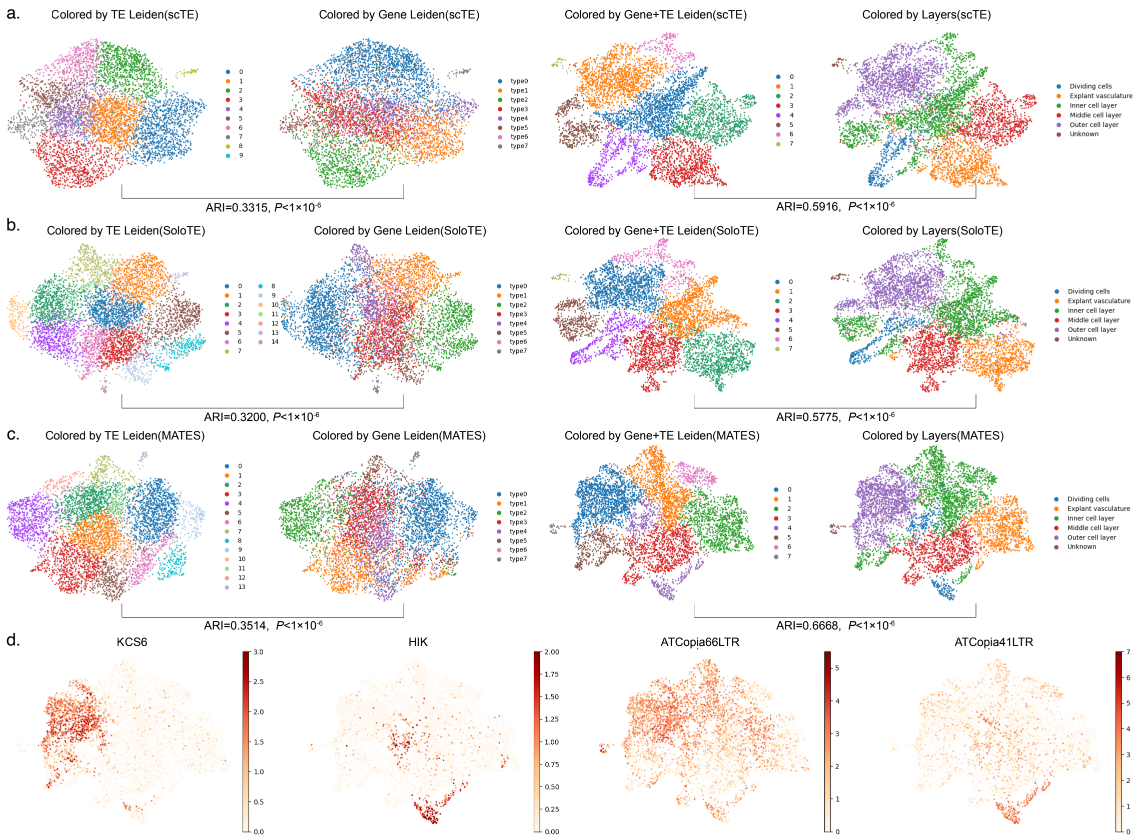


**Fig. S5 Cell Clustering Based on TE Expression and Gene Expression in Arabidopsis.** (a) scTE quantified TE expression. The left two panels show UMAP visualizations of cell clustering based on TE expression, colored by Leiden clusters and gene expression clusters (ARI: 0.3315). The right two panels show cell clustering based on Gene+TE expression, colored by Leiden clusters and layer labels(ARI:0.5916). (b) SoloTE quantified TE expression. The left two panels show UMAP visualizations of cell clustering based on TE expression, colored by Leiden clusters and gene expression clusters (ARI: 0.3200). The right two panels show cell clustering based on Gene+TE expression, colored by Leiden clusters and layer labels(ARI: 0.5775). (c) MATES quantified TE expression. The left two panels show UMAP visualizations of cell clustering based on TE expression, colored by Leiden clusters and gene expression clusters (ARI: 0.3514). The right two panels show cell clustering based on Gene+TE expression, colored by Leiden clusters and layer labels (ARI: 0.6686). (d) MATES identified Gene markers (left two panels) and TE markers (right two panels). The p-value of ARI is calculated against the random ARI score background built by using the block permutation strategy.

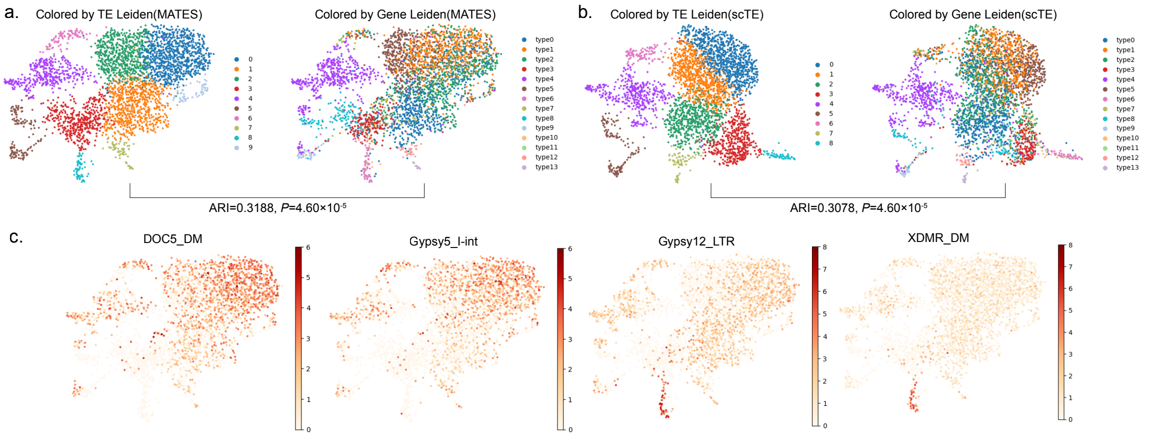


**Fig. S6 Cell Clustering Based on TE Expression and Gene Expression in Drosophila.** (a)MATES quantified TE expression. The left plot shows UMAP visualization of cell clustering based on TE expression with Leiden clusters, while the right plot shows the corresponding gene expression clusters with Leiden clusters. The ARI between the TE expression and gene expression clusters is 0.3188. (b)scTE quantified TE expression. The left plot shows UMAP visualization of cell clustering based on TE expression with Leiden clusters, while the right plot shows the corresponding gene expression clusters with Leiden clusters. The ARI between the TE expression and gene expression clusters is 0.3078. (c) MATES identified TE markers for different cell clusters. The p-value of ARI is calculated against the random ARI score background built by using the block permutation strategy.

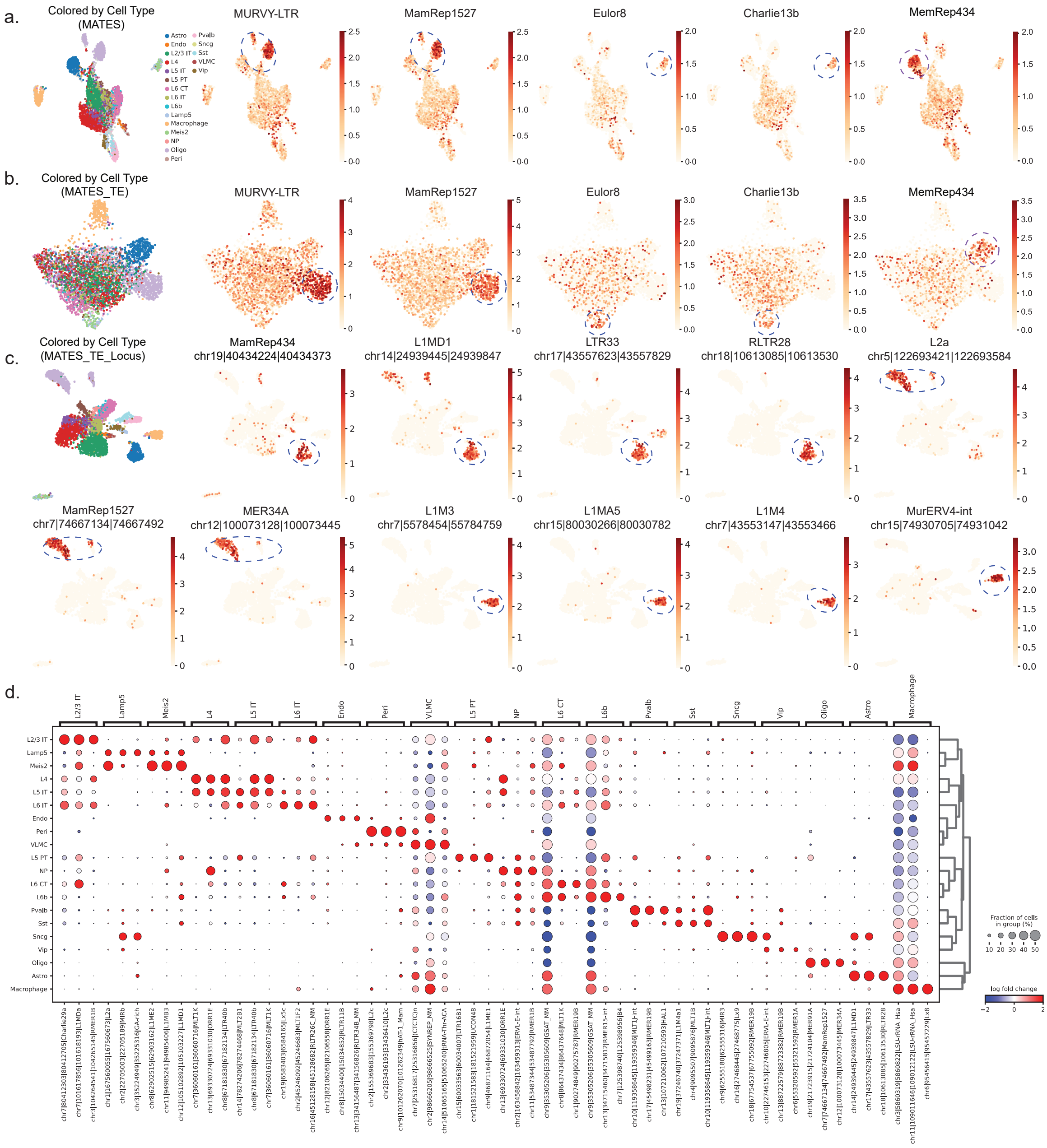

**Fig. S7 Extended Analysis of the 10X Single-Cell ATAC (scATAC-seq) Dataset from Adult Mouse Brain.** (a) UMAP visualization presents cell clustering using MATES quantification, which combines peak and comprehensive TE quantification. The UMAP is colored by cell type and TE markers, showcasing distinctiveness of Oligo cells marked by MURVY-LTR and MamRep1527, Meis2 cells by Eulor8 and Charlie 13b, and Astro cells by MamRep434. (b) Illustration of cell clustering exclusively based on TEs quantified by MATES, demonstrating the technique's capability for cell clustering and identification of characteristic TEs within the scATAC framework using only TE quantification. The UMAP follows the same color scheme by cell type and TE markers. (c) UMAP visualization showcasing cell clustering using locus-level TE expression, with colored by cell type and locus-specific TE markers: Astrocyte marked by MamRep434, L1MD1, LTR33, RLTR28; Oligo by L2a, MamRep1527, MER34A; and Macrophage by L1M3, L1MA5, L1M4, MurERV4-int. (d) A dot plot represents the marker TE locus for each cell type as identified by the MATES methodology.

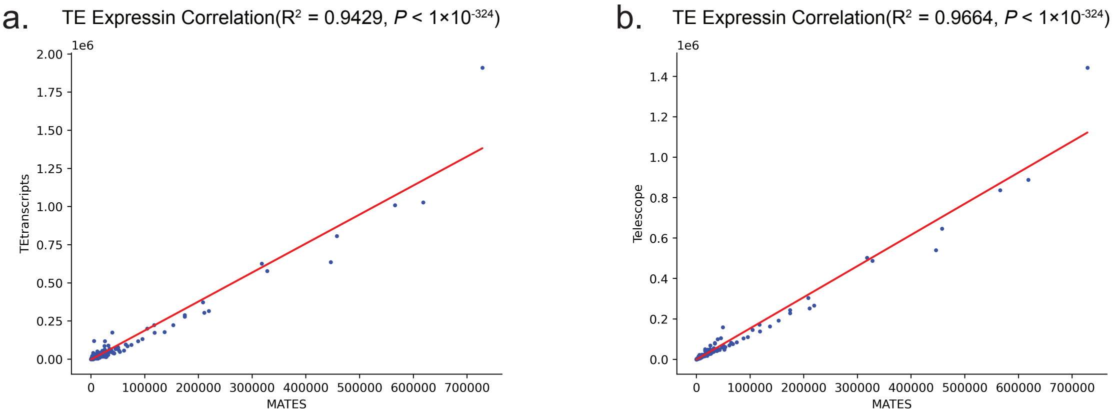


**Fig. S8 Comparison of Pseudo Bulk TE Expression Quantified by EM-based Quantification Methods and MATES.** (a) The scatter plot illustrates the correlation between TE expression levels quantified by TEtranscripts at the bulk level and by MATES at the single-cell level. To mimic bulk-level quantification for MATES, the average TE quantification across all cells was used. Each point represents the TE expression levels in pseudo-bulk data as quantified by the two methods. The Y-axis denotes the TE quantification obtained from bulk data using TEtranscripts. The X-axis represents the aggregated single-cell TE quantification by MATES. The red line indicates the linear regression fit, with an R-squared value of 0.9429, demonstrating a strong correlation between the quantification methods. Both methods have a p-value smaller than the precision limit ($P < 1 \times 10^{-324}$). The p-value of $R^2$ is calculated using the F-test.
(b) The scatter plot shows the correlation between TE expression levels quantified by Telescope at the bulk level and by MATES at the single-cell level. The Y-axis denotes the TE quantification obtained from bulk data using Telescope. The X-axis represents the aggregated single-cell TE quantification by MATES.The red line indicates the linear regression fit, with an R-squared value of 0.9664, demonstrating a strong correlation between the quantification methods. Both methods have a p-value smaller than the precision limit ($P < 1 \times 10^{-324}$). The p-value of $R^2$ is calculated using the F-test.

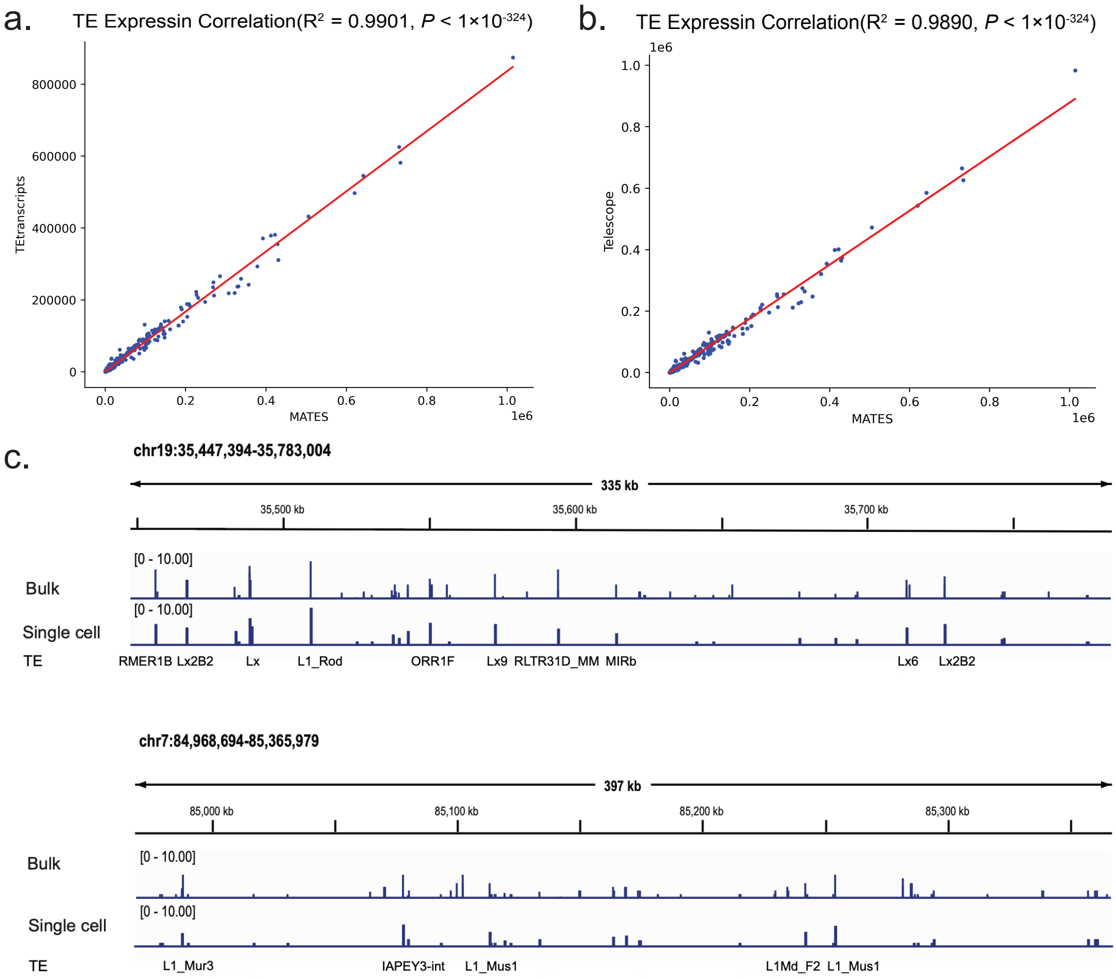


**Fig. S9 Comparison of MATES TE Quantification with Existing Bulk TE Quantification Methods on Pseudo-Bulk Data.** (a) The scatter plot illustrates the correlation between bulk-level TE expression levels quantified by TEtranscripts and the single-cell level TE quantification by MATES. Each point represents the TE expression levels in pseudo-bulk data as quantified by the two methods. The Y-axis denotes the TE quantification directly obtained from bulk data (pseudo-bulk BAM files) using TEtranscripts. The X-axis represents the aggregation of single-cell level TE quantification in all cells as quantified by MATES (the average of single-cell TE quantification for all cells by MATES). The red line indicates the linear regression fit, with an R-squared value of 0.9901, demonstrating a strong correlation between the quantification methods. Both methods have a p-value smaller than the precision limit ($P < 1 \times 10^{-324}$). The p-value of $R^2$ is calculated using the F-test.

(b) The scatter plot shows the comparison between Telescope and MATES. Each point represents the TE expression levels in pseudo-bulk data as quantified by the two methods. The Y-axis denotes the TE quantification directly obtained from bulk data (pseudo-bulk BAM files) using Telescope. The X-axis represents the aggregation of single-cell level TE quantification in all cells as quantified by MATES. The red line indicates the linear regression fit, with an R-squared value of 0.9890, demonstrating a strong correlation between the quantification methods. Both methods have a p-value smaller than the precision limit ($P < 1 \times 10^{-324}$). The p-value of $R^2$ is calculated using the F-test.

(c) IGV plots showing the agreement of single-cell (MATES) and bulk TE quantifications. This IGV plot contains two regions to show the agreement of MATES and bulk TE quantification results. The bulk and single-cell quantification results are shown in the plot, and the TEs of the overlapped peaks are annotated.

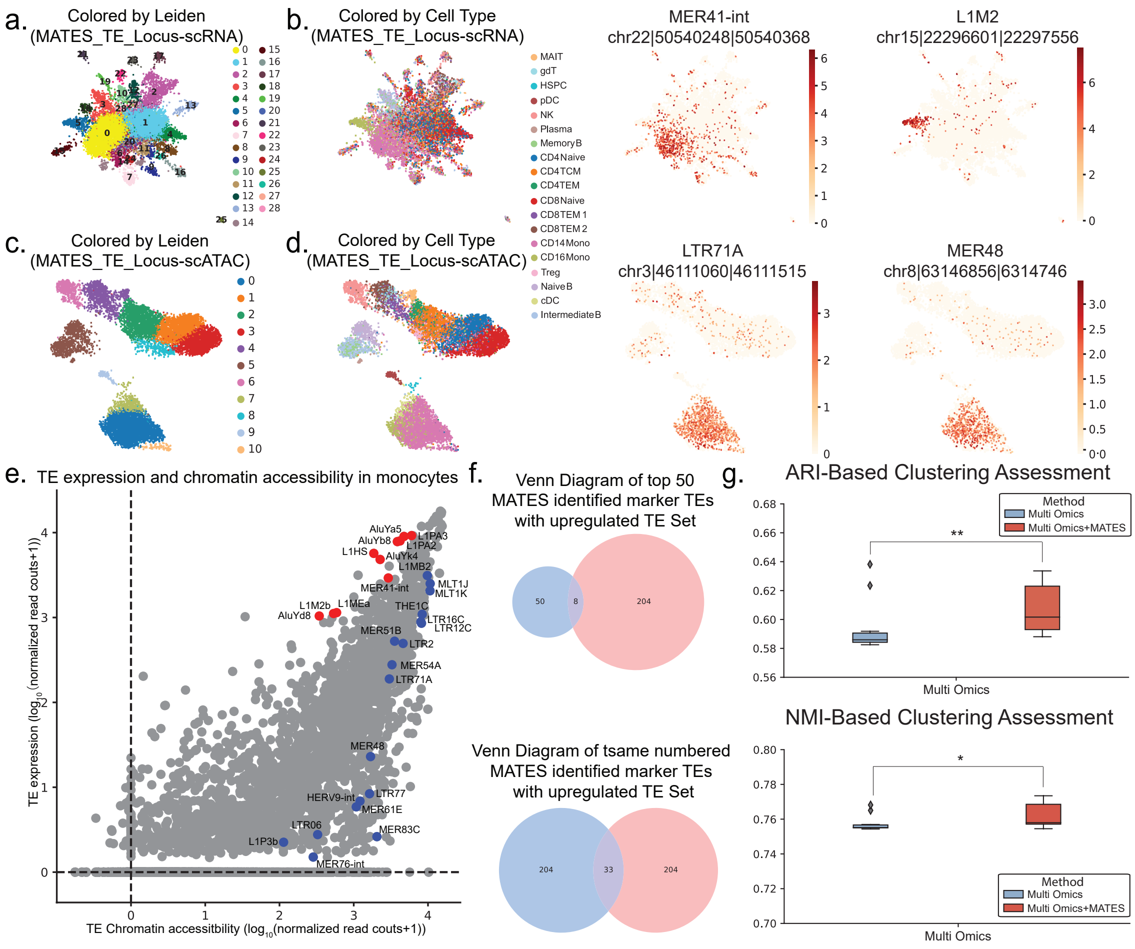


**Fig. S10 Supplementary MATES Analysis of the 10x Multiome PBMC Dataset.** (a-b) In UMAP visualization of cell clustering based on locus-level TE expression quantified by MATES using the scRNA modality, featuring locus TE biomarkers such as MER41-int and L1M2. Clusters are differentiated by the Leiden algorithm in (a) and by cell type and TE markers in (b). (c-d) UMAP visualization of cell clustering based on TE chromatin accessibility quantified by MATES in the scATAC modality, showcasing specific locus of TE biomarkers like LTR71A, and MER48. Clusters are colored by the Leiden in (c) and by cell type and TE markers in (d), illustrating the distinctive TE markers in each modality and the synergistic potential of MATES across both modalities. (e) A dot plot showcases the signature TEs for each cell type, highlighting the synergistic insights from both scATAC and scRNA modalities. (f) Venn diagram illustrating overlaps and distinctions among the top TE marker sets identified by MATES and upregulated TE sets post-influenza infection. (g) Comparison of traditional cell clustering techniques (represented by "Multi-omics" which combining scRNA_gene and scATAC_peak matrices) with the enhanced approach using "MATES" combined TE quantification. The p-values are calculated using the one-sided Student's t-test, with significance levels indicated as * for $P < 0.05$, ** for $P < 0.01$

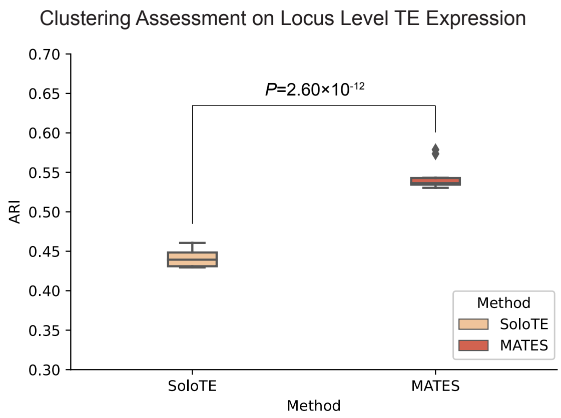


**Fig. S11 Locus-level benchmarking between MATES and SoloTE for TE expression quantification on 10X scRNA dataset.** The figure illustrates the ARI scores for cell clustering performance based on locus-level unique TE expression quantified by SoloTE and all TE expression quantified by MATES. The p-value was calculated using the one-sided Student's t-test.

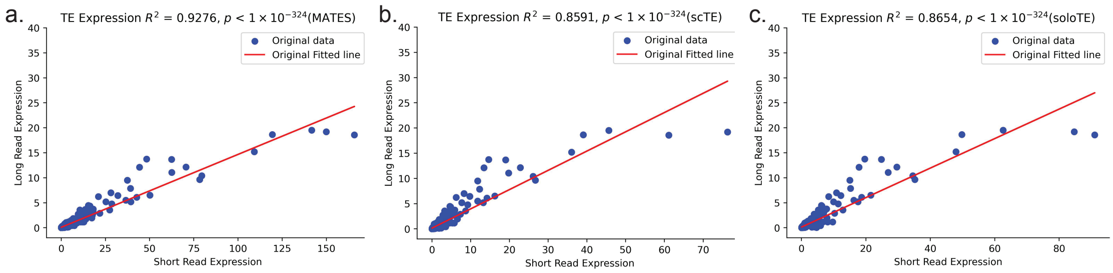


**Fig. S12 Additional Analysis for Validation:** (a)-(c) Pseudo-bulk analysis showing the correlation between long read TE expression and 10X short read expression quantified by different methods. Each plot compares the TE expression levels from Nanopore long read data to those quantified by (a)MATES, (b)scTE, and (c) SoloTE from the pseudo-bulk analysis of the short read data. The red line represents the linear regression fit. The p-values of $R^2$ were calculated using the F-test.

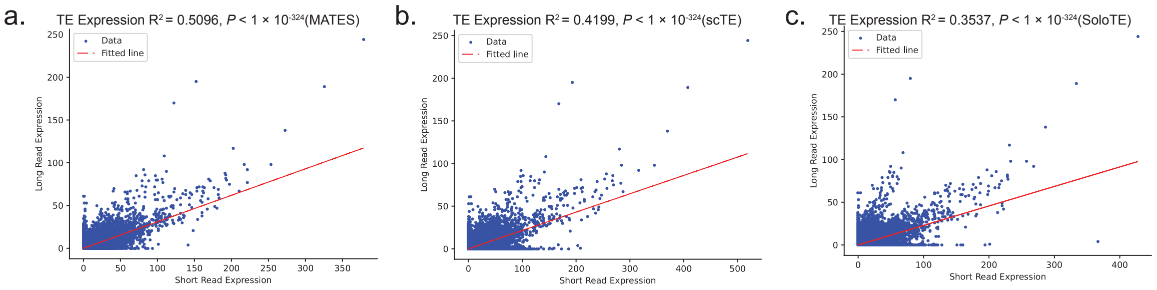


**Fig. S13 Validation on Pacbio long read data.** (a)-(c) Correlation of TE expression between long read data and TE expression quantified by different methods from paired 10X short read data. Each plot compares the TE expression levels from Pacbio long read data to those quantified by (a)MATES, (b)scTE, and (c) SoloTE from the paired short read data, respectively. The p-values of $R^2$ were calculated using the F-test.

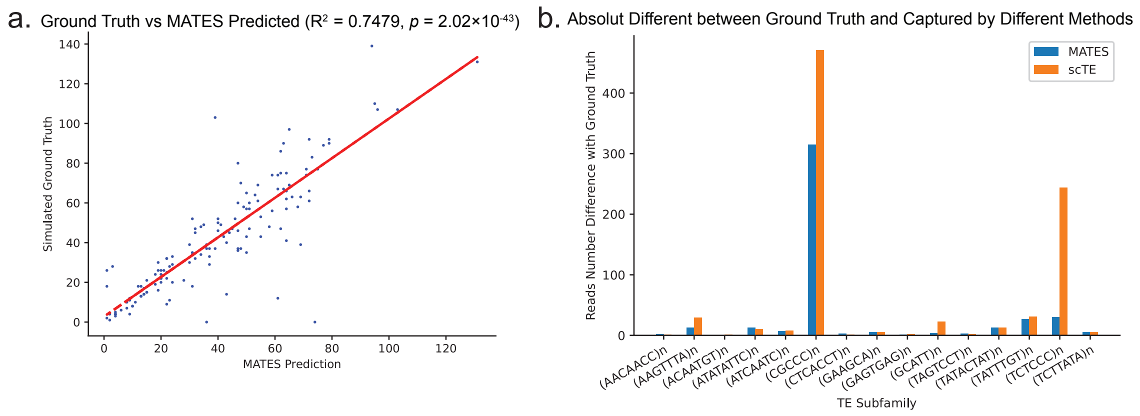


**Fig. S14 Evaluation of TE Expression Quantification for Simple Repeats using Simulation.** (a) Simulated Ground Truth vs. MATES Prediction ($R^2$= 0.7479). This scatter plot compares the simulated ground truth read numbers to MATES predictions, showing a high correlation. The p-value of $R^2$ was calculated using the F-test. (b) Captured Reads Difference with Ground Truth by Different Methods. This bar chart compares the read number differences between quantified and ground truth for MATES and scTE across various TE subfamilies. Shorter error bars indicate closer alignment to the ground truth.

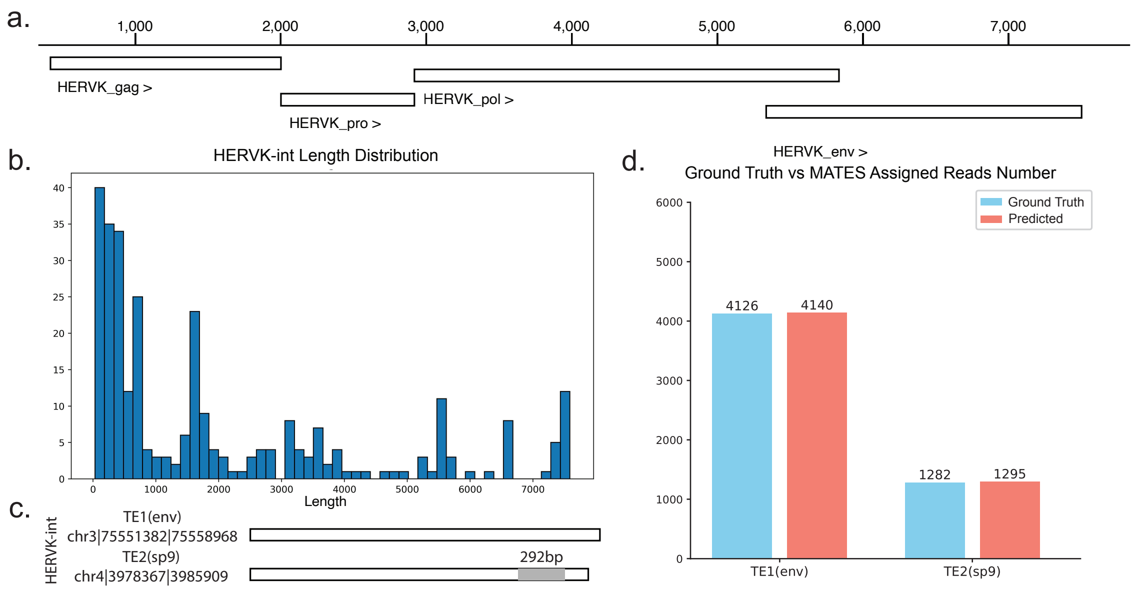


**Fig. S15 Simulation to Validate the Performance of MATES in Distinguishing Different HML2 isoforms.** (a) Semantic model of HML2 (HERVK). (b) The distribution of HERVK-int region lengths. (c) Selected full-length HERVK-int elements used for simulation. (d) MATES predicted TE isoform quantifications compared to the simulated ground truth.

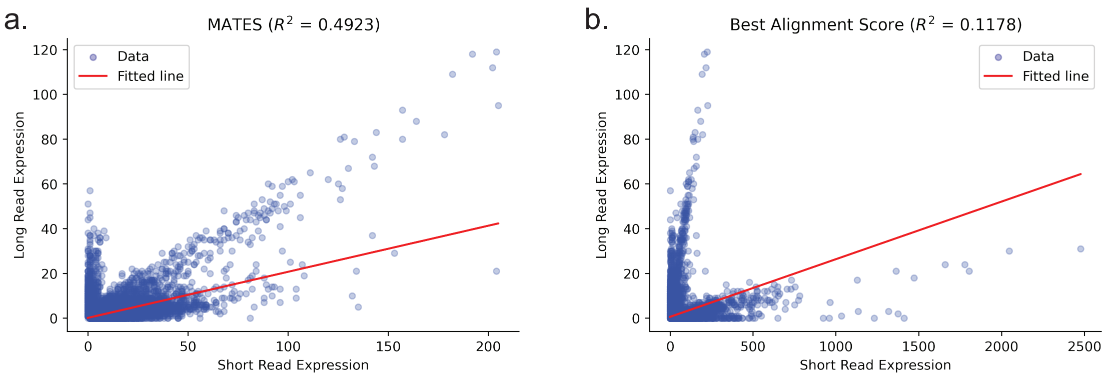


**Fig. S16 Benchmarking at the locus level.** Correlation assessment between locus-level TE expression quantified by (a) MATES ($R^2 = 0.4923, P < 1 \times 10^{-324}$) and (b) a random assignment strategy ($R^2 = 0.1178, P < 1 \times 10^{-324}$) on simulated short-read data, compared to the ground truth provided by nanopore long-read sequencing.

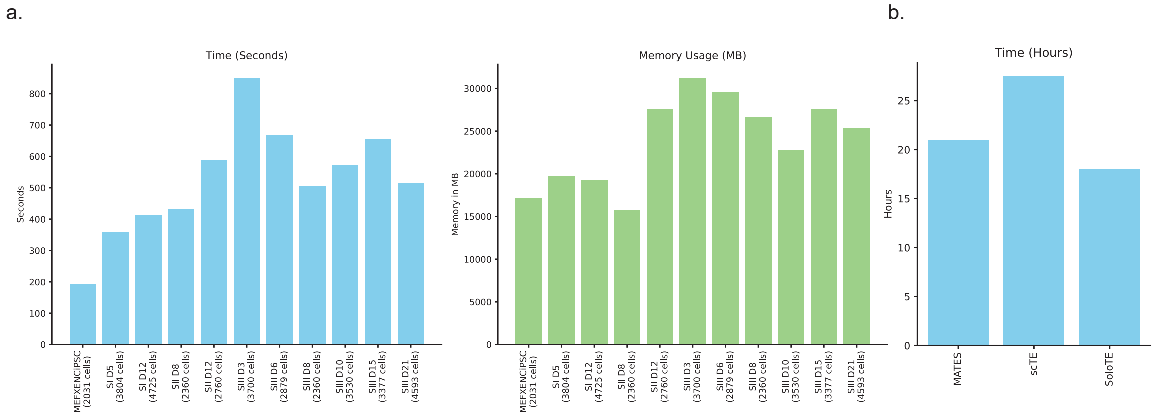


**Fig. S17 Analysis on Time and Memory Complexity** (a) Illustrates the time and memory consumption (in seconds and MB, respectively) of the model training and prediction process for samples at different reprogramming stages within the chemical reprogramming dataset. (b) Compares the overall time consumption (in hours) for quantifying the chemical reprogramming dataset using MATES, scTE, and SoloTE.

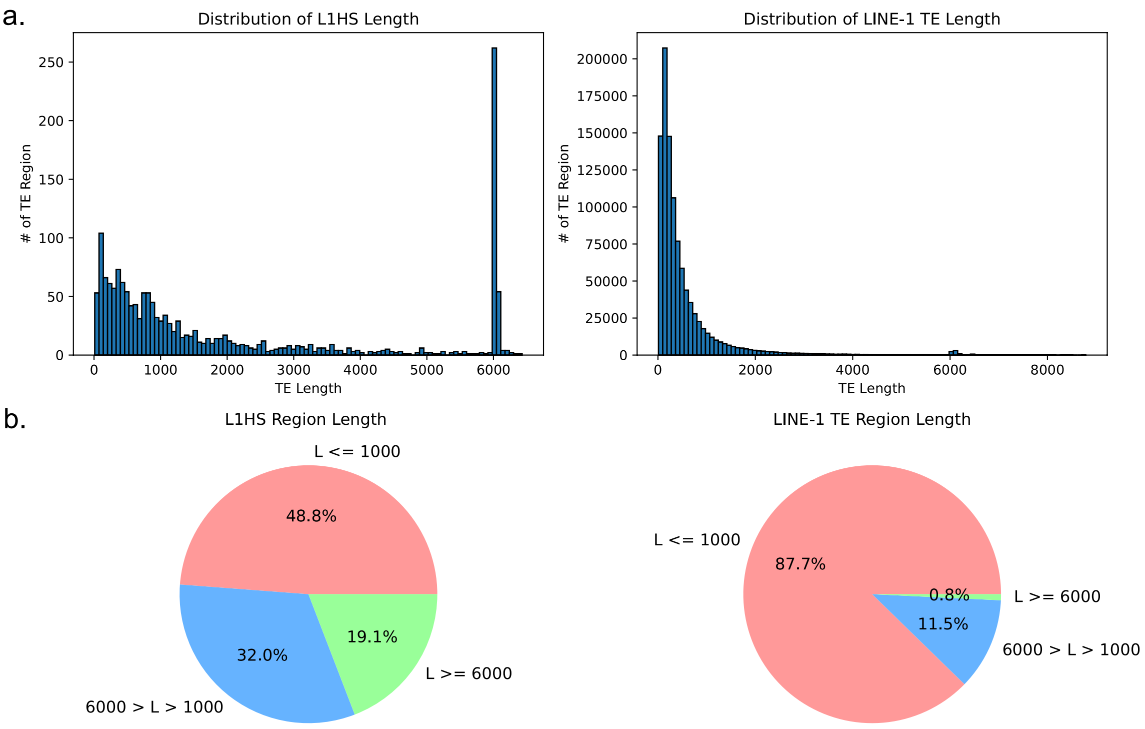


**Fig. S18 Distribution of TE region length for L1HS and LINE-1.** (a) Histograms display the distribution of TE region lengths for the L1HS and LINE-1 families. (b) A pie chart illustrates the proportions of different region lengths.

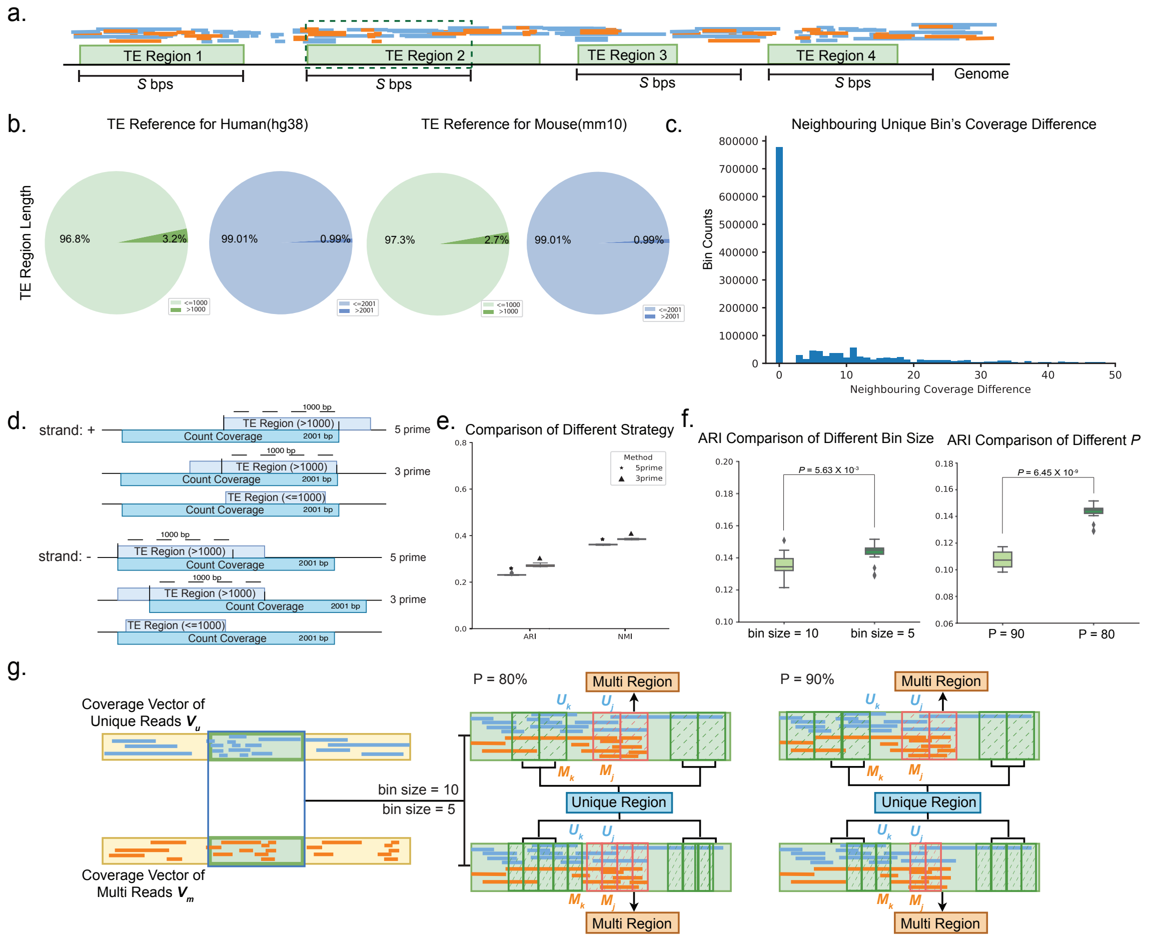

**Fig. S19 Explanation and Sections of MATES Hyperparameters.** (a) Illustrates the selection of hyperparameter $S$ and its impact on constructing the coverage vector. (b) Pie chart visualizations show that over $\sim$ 97% of TEs in both mouse and human genomes span less than 1000 bp, and over 99% of TEs in both mouse and human genomes span less than 2001 bp,suggesting that a 2001 bp window surrounding the TE is sufficient to capture the majority of TEs' local context in read alignment. (c) Analysis of coverage variation in TE regions, focusing on uniquely mapped reads, across adjacent bins. This analysis uses a subset of 150 cells randomly selected from 10X scRNA-seq data. (d) Schematic diagram of coverage vector construction in different scenarios. (e) Boxplots showing ARI and NMI scores for TE regions using 5' and 3' modes. Markers above the boxes indicate the different MATES modes. The performance using the 3' mode is generally better than the 5' mode in standard scRNA sequencing data. (f) Examines the effect of bin size and P value on cell clustering. In a 10x scRNA-seq dataset of chemical reprogramming, MATES indicates that a bin size of 5 generally provides better multi-mapping TE read quantification. However, performance across various bin sizes is relatively consistent. The role of $P$ in defining multi-mapping bins (M-bins) demonstrates that a lower $P$ value (e.g., $P$ = 80%) yields improved results, leading to default hyperparameters being set at bin size=5 and $P$ = 80%. (g)Discusses the impact of alterations in bin sizes and the percentage of multi-mapping reads ($P$) on both Multi-Region and Unique Region selection, affecting the base pairs covered by each region. Smaller bin sizes result in more stringent training sample selection, with a bin size of 5 recommended for samples with high multi-mapping rates, and larger sizes (e.g., bin_size=10) for those with lower rates. However, larger bin sizes may reduce performance due to less consistent read coverage continuity over extensive genomic regions.

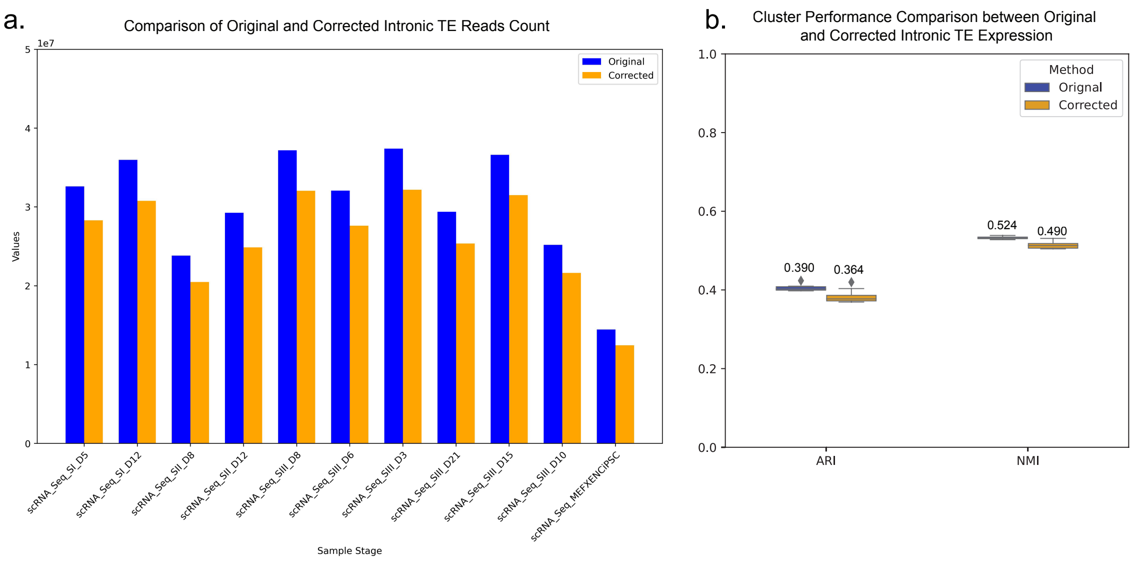


**Fig. S20 Comparison of Original and Corrected Intronic TE Reads Count and Clustering Performance.** (a) Comparison between the original number of reads aligned to intronic TEs and the MATES corrected intronic TE read counts across different sample stages. The blue bars represent the original read counts, while the orange bars represent the corrected read counts. (b) Clustering performance comparison based on original and MATES corrected intronic TE expression. The ARI and NMI scores are presented for both the original (blue) and corrected (yellow) methods. The median values are displayed above each box plot, and the p-values indicate the statistical significance of the differences between the original and corrected methods.
